# NeuroSCORE: A Genome-wide Omics-Based Model to Identify Candidate Disease Genes of the Central Nervous System

**DOI:** 10.1101/2021.02.04.429640

**Authors:** Kyle W. Davis, Colleen G. Bilancia, Megan Martin, Rena Vanzo, Megan Rimmasch, Yolanda Hom, Mohammed Uddin, Moises Serrano

## Abstract

To identify and prioritize candidate disease genes of the central nervous system (CNS) we created the Neurogenetic Systematic Correlation of Omics-Related Evidence (NeuroSCORE). We used five genome-wide metrics highly associated with neurological phenotypes to score 19,598 protein-coding genes. Genes scored one point per metric, resulting in a range of scores from 0-5. Approximately 13,000 genes were then paired with phenotype data from the Online Mendelian Inheritance in Man (OMIM) database. We used logistic regression to determine the odds ratio of each metric and compared genes scoring 1+ to cause a known CNS-related phenotype compared to genes that scored zero. We tested NeuroSCORE using microarray copy number variants (CNVs) in case-control cohorts, mouse model phenotype data, and gene ontology (GO) and pathway analyses. NeuroSCORE identified 8,296 genes scored ≥1, of which 1,580 are “high scoring” genes (scores ≥3). High scoring genes are significantly associated with CNS phenotypes (OR=5.5, *p*<2×10^−16^), enriched in case CNVs, and enriched in mouse ortholog genes associated with behavioral and nervous system abnormalities. GO and pathway analyses showed high scoring genes were enriched in chromatin remodeling, mRNA splicing, dendrite development, and neuron projection. OMIM has no phenotype for 1,062 high scoring genes (67%). Top scoring genes include *ANKRD17, CCAR1, CLASP1, DOCK9, EIF4G2, G3BP2, GRIA1, MAP4K4, MARK2, PCBP2, RNF145, SF1, SYNCRIP, TNPO2*, and *ZSWIM8*. NeuroSCORE identifies and prioritizes CNS-disease candidate genes, many not yet associated with any phenotype in OMIM. These findings can help direct future research and improve molecular diagnostics for individuals with neurological conditions.

## INTRODUCTION

A cholera epidemic swept across the globe in 1819 from India through the Middle East, Europe, and to America. London- and American-based physicians John Snow and Amariah Brigham both studied cholera and produced maps of the deaths in New York and London — Brigham’s in 1831 and Snow’s in 1855^1,2^. Both maps used different overlapping evidence, such as trade routes and drinking water systems, to illustrate a confluence of variables leading to new insights about cholera and, ultimately, public health remedies. Today, geneticists can take a similar approach, with different types of maps, to identify the genetic mechanism of diseases. With genome-wide, multi-omic analyses we can now overlay these datasets on the human genome and correlate these with phenotypes of the central nervous system (CNS) to identify candidate disease genes thereby improving diagnostics and, eventually, therapies.

Identifying disease or risk genes for conditions of the CNS has been a slow process, with current diagnostic rates for children with a broad range of neurological or developmental conditions ranging from 31% - 53%^3,4^ (undergoing multiple clinical tests) and approximately 32% in adults^5^. Diagnostic rates using exome and microarray vary within particular phenotypes, ranging from approximately 16% in autism spectrum disorder (ASD)^6^, 23% in corpus callosum anomalies^7^, and 42% in early-onset epileptic encephalopathies^8^. These conditions all likely have substantial unrecognized genetic contribution and a recent study of developmental disorders found that more than 1,000 additional genes are expected to contribute, either alone or in combination, to neurodevelopmental disorders^9^.

Identifying CNS-disease genes is complicated, as genetic CNS diseases are caused by multiple pathogenic mechanisms^4^, display multiple forms of inheritance, are characterized by allelic heterogeneity, reduced penetrance, pre/perinatal lethality, and variable expressivity, are difficult to study *in vivo*, have phenotypes that exist on a spectrum (e.g. ASD), have variable age of onset, have broad descriptions (e.g. “developmental delay”), and many genes characterized in the 1980s - 1990s have received disproportionate study leading to many unstudied genes, colloquially called the “ignorome”^10^.

Previous attempts to identify candidate disease genes have used multiple approaches, including statistical modeling for probability of causing a haploinsufficiency-related condition (pLI score^11^) or identifying genes with regional coding constraint^12^. Other systems rely on a particular type of variant observed in a control population (gnomAD’s observed/expected metrics^13^). Lastly, systems have been devised to search for disease-specific genes, such ForecASD^14^ and candidate ASD genes, or used specific data such as gene expression patterns in brain-coX^15^. These systems, while useful, are limited and no multi-omic system has yet been devised for CNS phenotypes.

As a diagnostic laboratory focused on neurological and developmental phenotypes, we sought to create a model that identified and prioritized candidate disease genes. We began with two foundational concepts, the first being “developmental brain dysfunction”, where the same condition can lead to a spectrum of central nervous system phenotypes (cognitive, motor, neurobehavioral, or neuroanatomical)^16,17^. Secondly, we used a multi-omics approach that accounts for different potential disease mechanisms and characteristics of genes that underlie known neurological conditions. Merging large scale omics databases supporting these two foundational concepts with a clinical database (i.e., the Online Mendelian Inheritance in Man (OMIM)), we created NeuroSCORE: the Neurogenic Systematic Correlation of Omics-Related evidence. We believe NeuroSCORE is the first multi-omic model to assess nearly all protein-coding genes and focused broadly on CNS phenotypes.

## METHODS

### Building NeuroSCORE Model

To build a comprehensive and clinically useful model, we chose publicly available, genome-wide databases with gene-specific data and combined them (Figure 1). As this analysis focused on protein-coding genes, we excluded non-coding genes, RNA genes, genes in mitochondrial DNA, and pseudogenes. We sought lines of evidence previously associated with neurodevelopmental or neurological phenotypes including loss of function constraint^11,13^, constrained coding regions^12^, *de novo* variation^18–25^, brain expression levels^26,27^, copy number variation^22,28^, and genes with exons that are both highly expressed in brain tissues and under purifying selection^29^. If a gene was identified by one of the following metrics, it received a score of one point (a categorical variable, yes vs. no). We began with seven preliminary gene metrics (possible independent variables) from which to build our model. Of note, two of these metrics have two levels, yielding nine total possible variables:

1. The gene’s upper bound score of the gnomAD observed/expected (o/e) loss-of-function metric was ≤0.34 (the preferred cutoff stated on the gnomAD site)^13^. Using gnomAD v2.1 (accessed May 2019), there were 2,832 genes identified by the gnomAD LOF gene metric.
2. The gene’s upper bound score of the gnomAD o/e missense variant metric was ≤0.34 (the preferred cutoff stated on the gnomAD site)^13^. The gnomAD MIS gene metric identified 119 genes.
3. The gene contained at least one area of regional constraint (critically constrained regions; CCRs) at or above the 95^th^ or 99^th^ percentile as described by Havrilla *et al.* (2019)^12^. The CCR 95 and CCR 99 gene metrics identified 7,049 and 1,444 genes, respectively.
4. The gene was enriched in the Coe, *et al*. (2014) case-control study of individuals with childhood developmental conditions and copy number variants (CNVs)^28^. We created two cutoff points for the Coe gene metrics based on two significance values: *p*≤0.01 (Coe 1=3,116 genes) or *p*≤0.02 (Coe 2=3,732 genes).
5. The median brain expression across the 13 brain tissues assessed by the Gene-Tissue Expression database v8 was ≥10 transcripts per million (“GTEx genes”)^30^. This cutoff was chosen based on the recommendion by the European Bioinformatics Institute (https://www.ebi.ac.uk/gxa/FAQ.html). The GTEx gene metric identified 6,069 genes.
6. The gene was enriched for *de novo* variants as reported in the *de novo* Database using the non-Simon Simplex Cohort data (assessed January 17^th^, 2020)^31^. Variants within protein-coding genes or the 3’ or 5’ untranslated regions from 13,166 trio or quartet exome/genome probands were collated from 31 unique studies for the following phenotypes: epilepsy, ASD, developmental delays, cerebral palsy, bipolar disorders 1 and 2, schizophrenia, early-onset Alzheimer and Parkinson disease, intellectual disability, neural tube defects, sporadic infantile spasm syndrome, and Tourette syndrome. We adopted a conservative approach to define a gene enriched with *de novo* variants if the genes contained ≥10 reported *de novo* variants. The *de novo* gene metric identified 487 genes.
7. The gene was identified by a previous exon indexing tool (“Index genes”) with exons expressed at or above the 90^th^ percentile in 388 post-mortem brain samples and below the 10^th^ percentile in mutational burden for rare (<5%) missense or loss-of-function variants in the 1,000 genomes database^29,32^. Although specific exons within a gene are identified with this tool, we scored the entire gene if ≥1 exon in the gene was identified. (Note: The cutoffs differ from those originally reported in the Uddin *et al.* (2016) paper as they are more stringent and are used by Lineagen, Inc. in interpretation of clinical testing.) The Index gene metric identified 4,646 genes.

**Figure 1:**
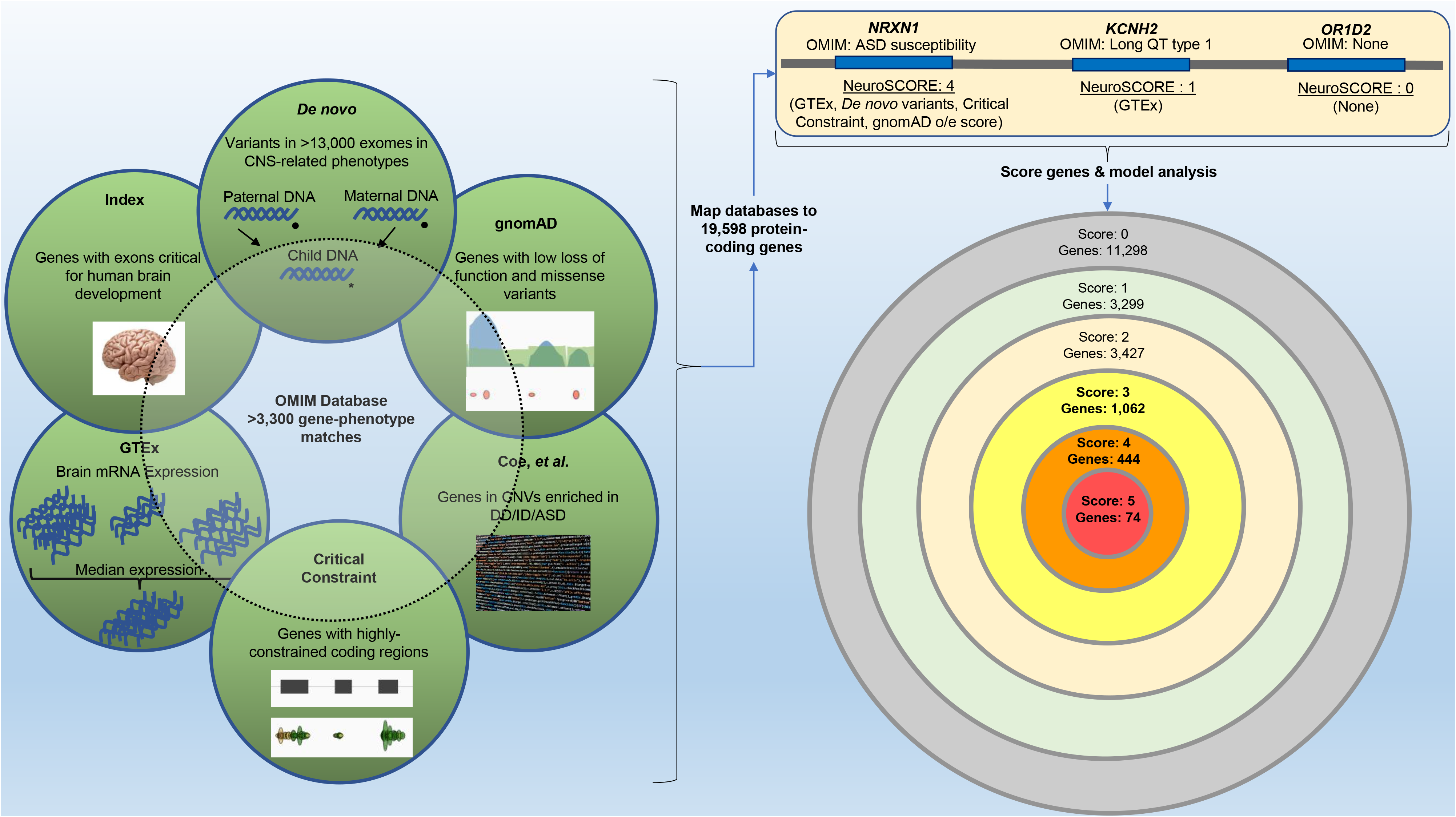
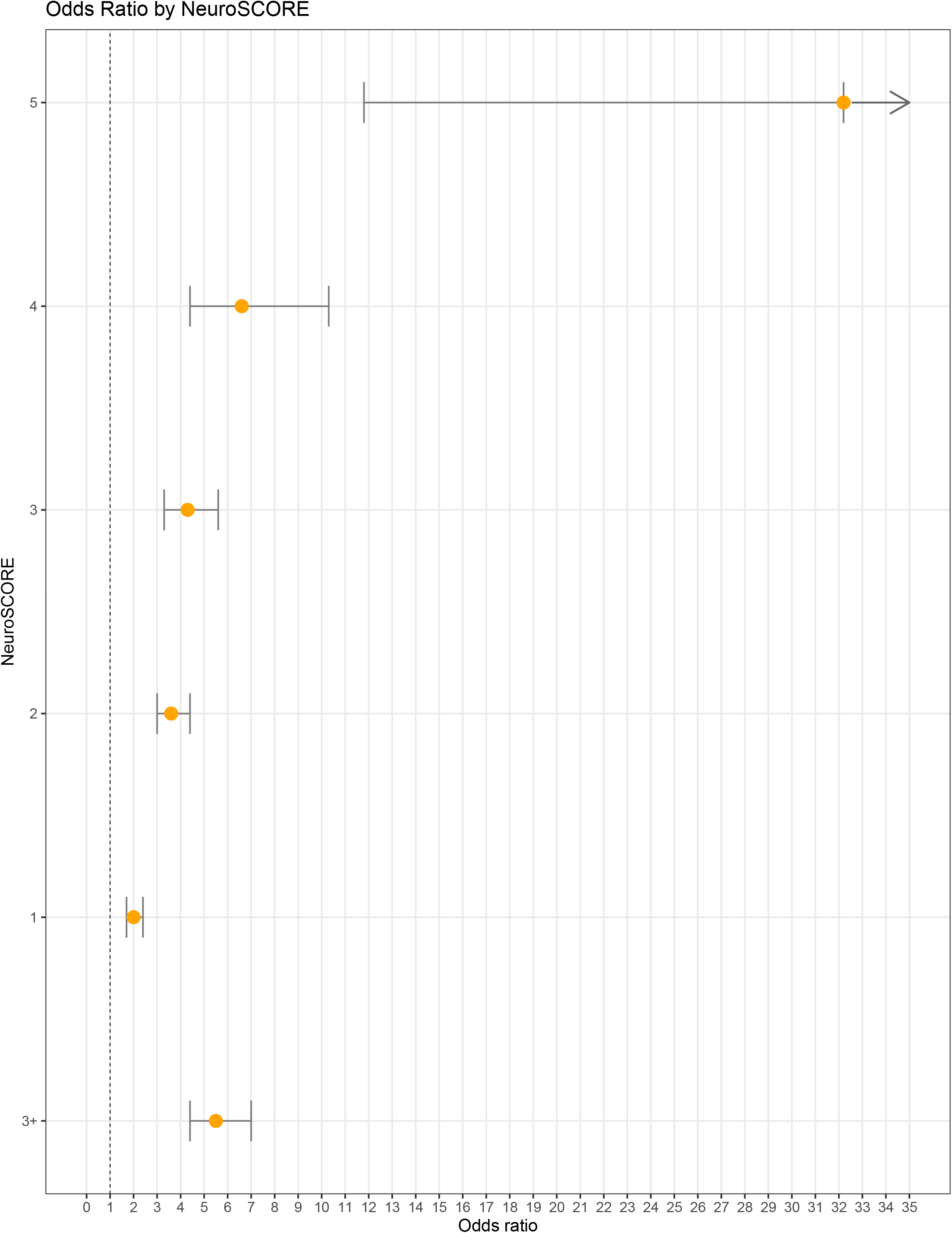
Schematic of NeuroSCORE Model Development.

Our outcome (independent) variable was defined as whether or not a gene was associated with a phenotype containing one or more CNS-related clinical features in OMIM and was also categorical (1 vs. 0, yes vs. no). We used a previous definition of the CNS as including only the brain and spinal cord^33^. We excluded phenotypes affecting the eye, retina, cochlea, and peripheral nervous system, or conditions that caused CNS involvement due to an external event (e.g., thromboembolism). Metabolic conditions and mitochondrial conditions caused by nuclear genes were included as having CNS phenotypes as the cellular dysfunction leading to symptoms originates within the cells of the CNS. Although the retina is derived from the CNS, phenotypes involving only the retina were excluded as these conditions are treated clinically as ophthalmological conditions. Two authors (KD and MR) reviewed the top 13,021 genes in OMIM (ranked by median brain expression) which included 1,822 genes with a phenotype including CNS-related clinical features and 1,513 genes without a CNS-related clinical features.

### Merging Databases and Data Fidelity

Due to genes having multiple historic names, we matched data between databases by both gene name and Ensemble ID. One author (KD) visually inspected all genes identified by the primary data sources and cross-referenced discrepant or missing data with external databases (e.g., HUGO) to ensure data was present. If a gene name was discrepant, the name was updated to the current HGNC-approved name. In total, our model assesses 19,162 genes, while data for 421 genes was not available from at least one gene metric.

### Statistical Analyses of the NeuroSCORE Model

We first assessed each of the seven gene metrics and their association with genes currently known to cause or contribute to CNS-related neurological phenotypes. We initially performed Pearson’s chi squared on each gene metric then included the variables significant at *p*<0.05 in a multiple logistic regression model. Nine total variables were assessed (seven metrics with the Coe and CCR gene metrics having two levels). Using SAS v9.4, we constructed a multiple logistic regression model with backward elimination to remove variables with high multicollinearity or those that were not associated with CNS-related clinical features at *p*<0.05. Wald testing was used to determine if each of the variables in the final model were significantly different from zero.

Using multiple logistic regression, we examined main effects and determined the odds ratios (ORs) for these metrics to be associated with genes known to cause CNS-related clinical features. We measured ORs for each variable in the final NeuroSCORE model as well as genes identified by multiple metrics (NeuroSCOREs 2-5). Both analyses used a comparison group of 4,723 genes that were not identified by any metric (NeuroSCORE of 0). This yielded 1,133 genes with a score of 0 that were linked to any known phenotype in OMIM (through June 2020). We performed a power analysis for these genes with a NeuroSCORE of 1 to determine the minimal detectable OR given our sample size. Setting *β*=0.95 and *α*=0.05 for this group of genes, the minimum OR we could detect was 1.4. All odds ratios were calculated using R using v.1.2.1335; power analyses were performed in R with the EpiR package v2.0.17.

### Evaluation of NeuroSCORE Model in Real-World Case and Control Cohorts

We used two published cohorts to evaluate NeuroSCORE. As exome analyses are often performed with priority to genes already known to be involved in genetic conditions, we used copy number variant (CNVs) from microarray data from individuals affected with neurological conditions and population control cohorts. This is because CNVs often contain multiple genes of known and unknown function and significance. We matched all genes in all included CNVs to their corresponding NeuroSCORE by gene name and visually inspected and corrected all discrepancies. CNVs with only non-scored genes (e.g., pseudogenes) were omitted from analysis.

The population control cohort was drawn from the Ontario Population Genomics Platform reporting on CNVs from 1,000 adults, providing 6,965 total CNVs^34^. After removing CNVs that did not affect at least one exon of one gene, the control cohort contained 1,862 gain CNVs and 2,547 loss CNVs. For the case comparison group, we began with a previously published cohort of 2,691 individuals with neurodevelopmental conditions including ASD, schizophrenia, attention deficit hyperactivity disorder (ADHD), and obsessive-compulsive disorder^35^. Almost half of this total cohort (46%, 1,230 / 2,691) was assessed for intellectual disability (ID), of which 14.9% (183/1,230) received the diagnosis and thus had ID combined with ASD, schizophrenia, or ADHD. We used CNVs consistently identified by multiple CNV calling algorithms (“stringent” CNVs) and were identified either as “rare” (<0.1% in a control population) or being deemed of possible clinical relevance (see Table 2 in Zarrei, *et al.* 2019). After removing CNVs from 17 individuals with aneuploidies, the final case data contained 1,357 gain CNVs and 835 loss CNVs. We included inheritance and clinical classification data when available.

For each CNV, we paired every gene with its respective NeuroSCORE, then generated a median and average NeuroSCORE for the CNV. To simplify these analyses, we converted NeuroSCOREs to percentages of the total possible points (1=20%, 2=40%, 3=60%, 4=80%, 5=100%). We used two-sided *t*-tests to compare differences in CNVs between cases and controls for CNV size, total gene content, and the average and median NeuroSCORE. We then used Pearson’s chi squared or Fisher’s exact test (if N≤5) to compare the distributions of the maximum scored gene within case and control CNVs. We performed sub-analyses using *t*-tests to explore differences in NeuroSCORE and scored genes by inheritance (inherited vs. *de novo*), gender (male vs. female proband), and clinical significance (common population variants, likely benign variants, variant of uncertain significance (VUS), and likely clinically significant/clinically significant). Finally, we performed linear regression analysis on classification and gene content by assigning increasing values to increasing pathogenicity and using the total count of genes in the CNV. Zarrei, *et al.* (2019) classified CNV pathogenicity, though we added a classification for common, “population variants” (CNVs observed at >1% in the cohorts). Classification was coded as 1=population variant, 2=likely benign, 3=VUS, 4=pathogenic/likely pathogenic.

### GO & Pathway Analyses

Gene ontology (GO) enrichment analysis was performed for the set of high scoring genes (identified by 3+ gene metrics in our final model; N=1,580)^36–38^. We performed analyses for biological processes, molecular function, and cellular component using Bonferroni correction for multiple testing. We chose this correction as it is the most conservative correction. If multiple related terms were within the top enriched GO terms, we included the more specific term and omitted the broader term. This analysis used databased from the Gene Ontology Consortium (http://geneontology.org/) using the March 23^rd^, 2020 release.

To map the relevant pathways in which the high scoring genes were primarily involved, gene enrichment analysis was performed using the gene overlap package of R, followed by Cytoscape analysis to trace the pathways involved and their connectivity. The false decision rate and *p*-value cut off was 0.01 and 0.001, respectively. Kyoto Encyclopedia of Genes and Genomes (KEGG) and Gene Ontology (GO) database were used for both gene enrichment and Cytoscape analysis. Then, the network was built using the enrichment map and the auto annotate Cytoscape application. The node color represents the *p*-value (the darker the shade, the lower the *p*-value) and size of the node represents increasing odds ratio.

### Evaluation of NeuroSCORE Model in Mouse Model Data

We used the high-level phenotypic data provided by the Jackson Laboratory’s Mouse Genome Informatics database to further assess our model (accessed July 6^th^, 2020; http://www.informatics.jax.org)^39^. We downloaded all annotated genes and matched the mouse and human gene using the unique MGI number via HGNC database. We excluded genes where the human homolog of the mouse gene included two or more unique human genes (e.g., the mouse gene *Rln1* is a homolog of both human *RLN1* and *RLN2*). We also removed multiple mouse genes that matched the same human ortholog (e.g., mouse genes *SCD1*, *SCD2*, *SCD3*, and *SCD4* are homologues of human *SCD*). Lastly, we removed genes where no mouse phenotype information was available, as a lack of high-level phenotype information does not mean variants of a gene cannot cause a phenotype. In total, we removed: 70 genes with multiple mouse homologues matching to a single human gene, 45 human genes that did not have a mouse ortholog, and 4,139 genes without phenotype data. The total number of genes included for analysis with phenotype data was 8,149. Of note, this data was not used to develop or refine the NeuroSCORE model.

### Data Availability

The data that support the findings of this study are available from Bionano Genomics, Inc., but restrictions apply to the availability of these data, which were used under license for the current study, and so are not publicly available. Data, however, may be available from the authors upon reasonable request and with permission of Bionano Genomics, Inc.

## RESULTS

### Constructing and Assessing the NeuroSCORE Model

Using Pearson’s chi squared, we determined if any of the nine gene metrics (dependent variables) was correlated with our outcome measure of genes currently associated with a CNS-related condition. Eight of the nine variables were significantly associated with currently known CNS-related disease genes including: GTEx genes, *de novo* genes, critically constrained genes at both the 99^th^ and 95^th^ percentiles, gnomAD LOF and MIS genes, Index genes, and Coe 1 genes (all *p*<.05). The Coe 2 genes were not correlated (*p*=.10). Using these eight metrics, we then constructed a multiple logistic regression model. Main effects of logistic regression results indicated that five of the eight variables were significantly and positively associated with odds ratios (ORs) above 1.0 for the outcome measure of CNS-related disease genes (Table 1). This model with five variables—*de novo* genes, Index genes, critically constrained genes at the 99^th^ percentile, GTEx genes, gnomAD LOF genes—became our final NeuroSCORE model.

**Table 1:** Gene Metric Association of Genes with CNS-Related Clinical Features

| Gene Metric | Pearson's $\chi^2$ | | Logistic Regression | | | Wald Test | |
| --- | --- | --- | --- | --- | --- | --- | --- |
| | $\chi^2$ | $p$ | OR | 95% CI | $p$ | $F$ | $p$ |
| *De novo* | 67.0 | <1E<sup>-4</sup> | 2.2 | 1.7 – 3.0 | <1E<sup>-4</sup> | 27.5 | <1E<sup>-4</sup> |
| Index | 171.1 | <1E<sup>-4</sup> | 1.9 | 1.5 – 2.3 | <1E<sup>-4</sup> | 39.4 | <1E<sup>-4</sup> |
| CCR 99 | 82.9 | <1E<sup>-4</sup> | 1.8 | 1.4 – 2.3 | <1E<sup>-4</sup> | 24.3 | <1E<sup>-4</sup> |
| GTE<sub>x</sub> | 154.0 | <1E<sup>-4</sup> | 1.7 | 1.4 – 2.0 | <1E<sup>-4</sup> | 30.7 | <1E<sup>-4</sup> |
| gnomAD LOF | 55.5 | <1E<sup>-4</sup> | 1.4 | 1.1 – 1.6 | 5E<sup>-4</sup> | 12.1 | 5E<sup>-4</sup> |
| CCR 95 | 19.8 | <1E<sup>-4</sup> | 0.9 | 0.8 – 1.1 | 0.4 | NA | NA |
| gnomAD MIS | 20.2 | <1E<sup>-4</sup> | 1.8 | 0.8 – 4.5 | 0.2 | NA | NA |
| Coe 1 | 4.2 | .04 | 1.2 | 0.9 – 1.4 | .10 | NA | NA |
| Coe 2 | 2.7 | .10 | NA | NA | NA | NA | NA |

OR: odds ratio; 95% CI: 95% confidence interval; CCR 95 or 99: genes with $\geq 1$ critically constrained coding region at the 95<sup>th</sup> or 99<sup>th</sup> percentiles; GTE<sub>x</sub>: gene-tissue expression database; gnomAD LOF and MIS: gnomAD genes with upper bound of loss-of-function or missense observed/expected metric <0.34; NA: not applicable.

NeuroSCORE creates different scoring levels for genes from 1-5 points. We then used logistic regression to investigate the relative enrichment of currently known CNS-related disease genes within each scoring level compared to genes that scored zero and calculated the OR for genes identified at each scoring level (Table 2 and Figure 2). Odds ratios increased with each increase in NeuroSCORE, ranging from 2.0 – 32.2 (all *p*<5E^−8^). Next, we calculated the OR for genes that had a majority of the NeuroSCORE points (≥3), which we considered to be “high scoring” genes (Table 2). As this set of high scoring genes is significantly associated with CNS-related disease genes, we focused the remaining analyses on the high scoring genes (N=1,580).

**FIgure 2:**
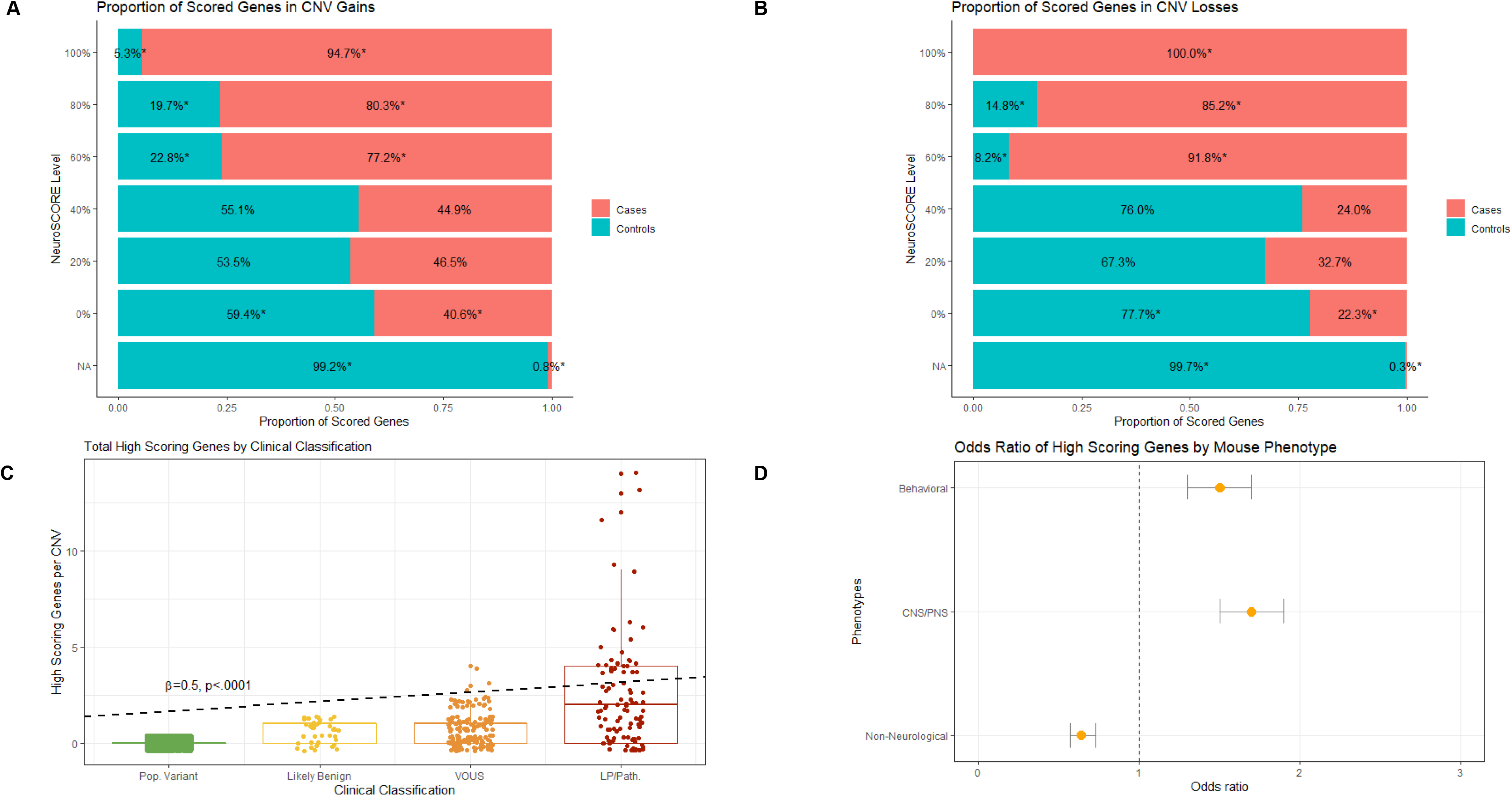
A:The relative proportion of genes at each NeuroSCORE level in loss CNVs, \**p*≤.002; **B:** relative proportion of genes at each NeuroSCORE level in gain CNVs; **C:** Box plots of high scoring genes in CNVs by clinical significance; **D:** High scoring genes are enriched in in behavioral (OR=1.5, 95% CI: 1.3-1.7, \**p*<.001) and central and peripheral nervous systems (OR=1.7, 95% CI: 1.5-1.9, \**p*<.001) mouse phenotypes and depleted in non-neurological phenotypes (OR=.65, 95% CI: .57-.73, *p*<.001).

**Table 2:** NeuroSCORE Model Shows Increasing Odds Ratios with Increasing Point Totals

| Point Total | OR | 95% CI | *p* | Total Genes<br>(Genes without OMIM Phenotype) |
| --- | --- | --- | --- | --- |
| 5 of 5 | 32.2 | 11.8 – 132.7 | 5.7E<sup>-9</sup> | 74 (15) |
| 4 of 5 | 6.6 | 4.4 – 10.3 | 2.0E<sup>-16</sup> | 444 (302) |
| 3 of 5 | 4.3 | 3.3 – 5.6 | 2.0E<sup>-16</sup> | 1,062 (745) |
| 2 of 5 | 3.6 | 3.0 – 4.4 | 2.0E<sup>-16</sup> | 3,423 (2,560) |
| 1 of 5 | 2.0 | 1.7 – 2.4 | 1.1E<sup>-14</sup> | 3,293 (2,431) |
| ≥3 of 5 | 5.5 | 4.4 – 7.0 | 2.0E<sup>-16</sup> | 1,580 (1,062) |

OR: odds ratio; 95% CI: 95% confidence interval; high scoring genes are genes identified by ≥3 gene sets; OMIM data current as of December 31<sup>st</sup>, 2020

### Case-Control Analyses in a Neurodevelopmentally Affected Cohort

We applied NeuroSCORE to copy number variant (CNV) data derived from microarray testing in both a population control^34^ and a neurodevelopmentally affected cohort.^35^ We compared average CNV size, total genes involved, total number of genes identified by NeuroSCORE, and median and average NueroSCORE of the CNV (*p-*values used Bonferroni correction of *p*<.005 corresponding to five statistical tests for two classes of CNVs). For easier interpretation of the following analyses, NeuroSCOREs were converted from total points to a percentage of total points (0=0%, 1=20%, 2=40%, 3=60%, 4=80%, and 5=100%).

Using *t-*tests, we found the average size of both loss and gain CNVs was significantly larger in cases than controls (losses: 344 kilobases (kb) vs. 104 kb, *p*=3E^−8^; gains: 419 kb vs. 244 kb, *p*=5E^−8^). Within gain CNVs, the average number of genes was similar between cases and controls (4.1 v. 3.6, *p*=.1), while there were more genes on average within loss CNVs for cases versus controls (3.2 vs. 2.4, *p*=.001). The similar total number of genes within case and control CNVs suggests that *specific* genes within case CNVs, rather than total genes, are more likely to contribute the clinical neurodevelopmental phenotype.

To assess potential differences in gene content, we next analyzed median and average CNV NeuroSCORE and the distribution of scored genes between case and control CNVs. Using *t*-tests, we found that case CNVs had higher median and average NeuroSCORE (median: losses: 13.4% vs. 6.7%, *p*=2E^−16^; gains: 14.9% vs. 8.5%, *p*=2E^−16^; average: losses: 14.2% vs 7.6%, *p*=2E^−16^; gains: 16.4% vs. 10.5%, *p*=2E^−16^). To assess the distribution of scored genes within CNVs, we used chi squared analyses using the highest scoring gene in a CNV at each different scoring levels (100%, 80%, 60%, 40%, 20%, 0%, and no score/NA). Using a Bonferroni corrected *p*-value for 14 tests of *p*<.004 (7 scoring levels, two classes of CNVs), we found that case CNVs were significantly enriched for the high scoring genes in both loss and gain CNVs while controls were enriched for genes that achieved no score (0%) or were not scored/NA (Table 3 and Figure 3A and B). In case CNVs, 16.2% of losses and 21.9% of gains contained at least one high scoring gene compared to 0.7% and 5.3% of controls, respectively. Logistic regression showed a significantly increased OR for case CNVs containing one or more high scoring genes compared to controls (OR = 9.3, 95% CI: 7.4 – 11.8, *p*=2e^−16^).

**Table 3:** Distribution of the Highest Scored Gene Within Case & Control CNVs

| Point<br>Total | Genes in Loss CNVs |  |  | Genes in Gain CNVs |  |  |
| --- | --- | --- | --- | --- | --- | --- |
|  | Cases<br>N (%) | Controls<br>N (%) | *p* | Cases<br>N (%) | Controls<br>N (%) | *p* |
| 5 | 5 (0.6) | 0 (0) | $1.6E^{-3}$ | 17 (1.3) | 1 (0.1) | $2E^{-5}$ |
| 4 | 53 (6.5) | 9 (0.4) | $2E^{-16}$ | 81 (6.1) | 25 (1.7) | $8E^{-10}$ |
| 3 | 60 (7.4) | 7 (0.3) | $2E^{-16}$ | 165 (12.5) | 52 (3.5) | $2E^{-16}$ |
| 2 | 115 (14.1) | 365 (16.7) | .09 | 285 (21.6) | 354 (24.0) | .13 |
| 1 | 114 (17.7) | 307 (14.0) | .01 | 260 (19.7) | 300 (20.3) | .68 |
| 0 | 436 (53.6) | 1501 (68.6) | $3E^{-14}$ | 514 (38.9) | 746 (50.5) | $7E^{-10}$ |
| NA | 1 (0.1) | 358 (14.1) | $2E^{-16}$ | 3 (0.2) | 384 (20.6) | $2E^{-16}$ |

NA are genes that could not be scored (e.g., pseudogenes); total case CNVs $N_{\text{LOSSES}}=813$ , $N_{\text{GAINS}}=1,322$ , total control CNVs $N_{\text{LOSSES}}=2,547$ , $N_{\text{GAINS}}=1,862$ ; Fisher's Exact test used for analyses when $N \leq 5$ ; Chi squared tests used for analyses when $N > 5$ ; significance set at $p \leq 4E^{-3}$ after Bonferroni correction for 14 tests

**Figure 3:**
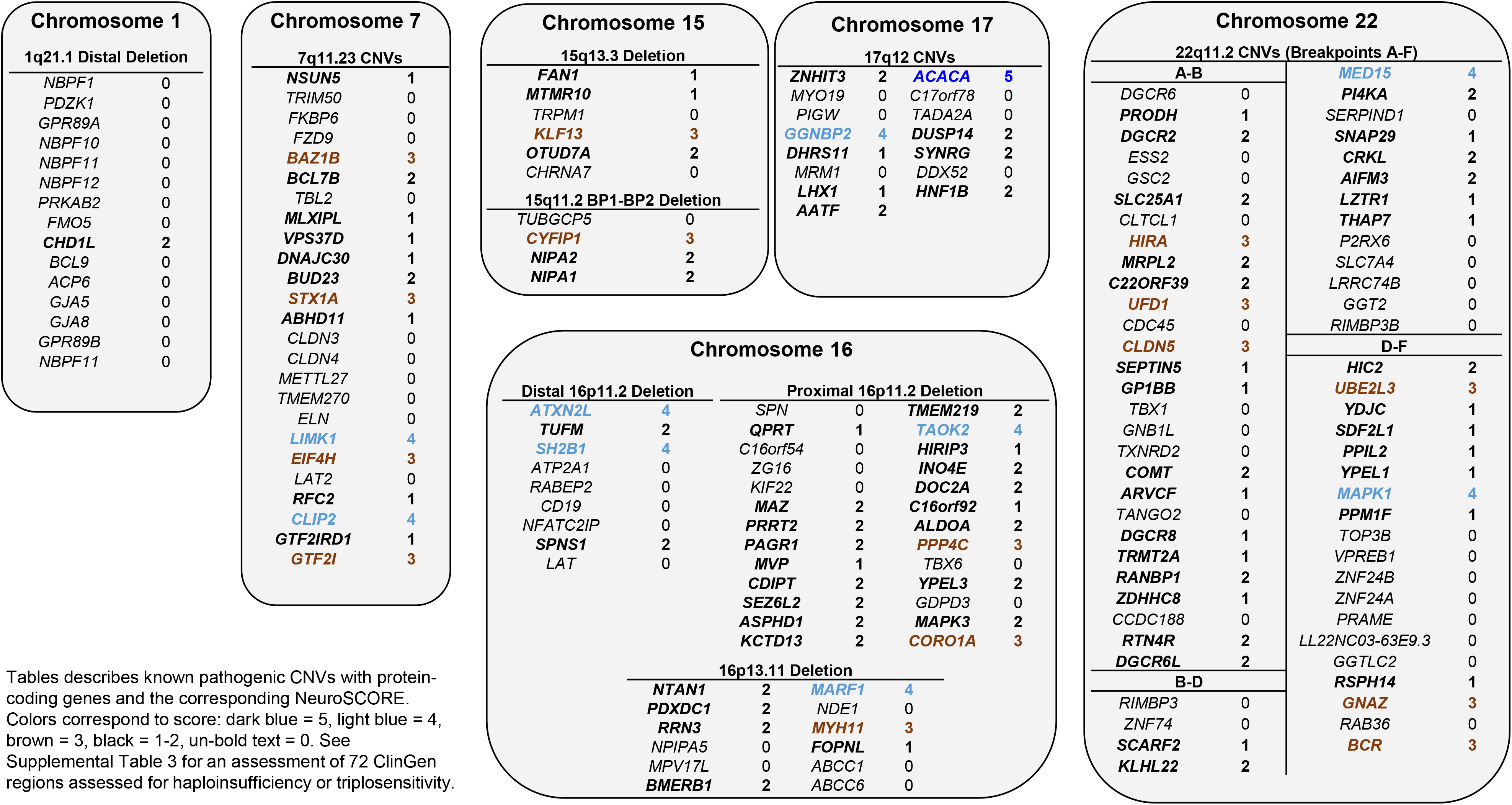
Common Neurodevelomental Microdeletion/Duplication Syndromes with Gene Level NeuroSCORE.

We next performed sub-analyses of case-control CNV by gender (N_MALES_=1,724, N_FEMALES_=468), inheritance (N_PATERNAL_=35, N_MATERNAL_=43, N_*de novo*_=33), and clinical classification (N_POPULATION_=2,195, N_LIKELY BENIGN_=39, N_VOUS_=148, N_CLIN. SIG_=96; Figure 3C). Previous work has shown that the gender bias in neurodevelopmental conditions is partly due to the burden of CNVs or sequence variants^40^. Given this, we questioned if affected females had CNVs with higher NeuroSCOREs than affected males. Analyzing case CNVs by gender showed no differences for average NeuroSCORE (15.1% vs. 15.7%), median NeuroSCORE (13.7% vs. 14.6%), or rates of high scoring genes (0.4 vs. 0.3). Similarly, we did not find differences in CNVs by inheritance (inherited vs. *de novo*) for average NeuroSCORE (20.6% vs. 26.1%), median NeuroSCORE (17.3% vs. 25.2%), or rates of high scoring genes (1.8 vs. 0.7). However, using linear regression and controlling for CNV size we found that CNVs with increasing pathogenicity showed an increase in the total number of high scoring genes (*β*=0.5; *p*=2E^−16^) and lower scoring genes (*β*=0.02; *p*=.02), while non-scored genes decreased slightly (*β*= −0.05; *p*=2.5E^−8,^ Figure 3C). As expected, linear regression also showed median CNV NeuroSCORE increased with increasing classification as well (*β*=6.7%; *p*<2 E^−16^). These data show high scoring genes appear to be an important component in CNVs identified in this clinically affected cohort.

### High Scoring Genes and Mouse Model Organism Data

We next applied NeuroSCORE to the 27 high-level phenotypes of mouse model data curated by Jackson Labs Mouse Genome Informatics database^39^. Among genes with high-level experimental phenotype data (N=8,149), total phenotypes per gene ranged from 1-26 with an average of 5.99 phenotypes (st. dev. = 4.55). Using chi squared analysis between high scoring gene orthologs and the presence of any of the 27 phenotypes (Bonferroni corrected for 27 tests, *p*<0.002), we found high scoring gene orthologs were significantly enriched in seven phenotypes including: mortality and aging, embryonic abnormalities, central and peripheral nervous system abnormalities, growth and congenital anomalies, abnormalities of learning and behavior, abnormalities of cellular proliferation, differentiation, and apoptosis, and abnormal muscle development (all *p*≤2.9E^−5^). Using logistic regression, we found high scoring genes were significantly more likely to be associated with mouse ortholog genes that cause the behavioral phenotypes (OR=1.5, 95% CI:1.3 – 1.7, *p=*5.9E^−10^) and central/peripheral nervous system phenotypes (OR=1.7, 95% CI: 1.5 – 1.9, *p*=4.7E^−16^), while they were significantly depleted in non-neurological phenotypes (OR=.64, 95% CI: .57 - .73, *p=*3.8E^−12^; see Figure 3D).

### GO Enrichment and Pathway Analyses

We performed GO Analyses on the set of 1,580 high scoring genes to determine if enrichment of key biological processes, cellular components, and molecular functions occurred within this set. Enriched terms in biological processes include positive regulation of protein localization to Cajal body (GO:1904871), axo-dendritic protein transport (GO:0099640), and alternative mRNA splicing, via spliceosome (GO:0000380). Within cellular components, key terms include the nBAF complex (GO:0071565) and the NuRD complex (GO:0016581), which are involved in chromatin remodeling. For the molecular functioning area, terms include binding activity such as protein kinase A catalytic subunit binding (GO:0034236), microtubule plus-end binding (GO:0051010), and pre-mRNA binding (GO:0036002). See Supplemental Table 1 for a list of all of the top five enriched, unique GO annotation terms and high scoring genes for each of the three areas.

From our pathway analyses, we analyzed approximately 500 GO terms with the lowest false discovery rate values (all q-values ≤1.3E^−22^) by inspecting the relationships using the AmiGO visualization tool (accessed December 10^th^, 2020). Within the developmental pathways, the following were enriched: regulation of dendrite development (GO:0050773), regulation of morphogenesis involved in differentiation (GO:0010769), positive regulation of neurogenesis (GO:0050769), and positive regulation of neuron projection development (GO:0010976). Within the metabolic and enzymatic pathways, the following were enriched: regulation of mRNA stability (GO:0043488), regulation of mRNA splicing via the spliceosome (GO:0048024), and catalytic step two of the spliceosome (GO:0071013). Top selected terms and associated genes are presented in Supplemental Table 2 and the corresponding figure. The GO and pathway analyses further support that NeuroSCORE identifies genes in important neurological and developmental processes.

### Landscape of High Scoring Genes in Pathogenic CNVs

Given pathogenic CNVs underlie a significant proportion of many neurological disorders, we applied NeuroSCORE to a set of common pathogenic CNVs as described in >10,000 individuals with neurological or neurodevelopmental phenotypes^41^. All CNVs had at least one scored gene and all but the 1q21.1 distal deletion CNV had at least one high scoring gene (Figure 4). We next applied NeuroSCORE to the 72 CNV regions with a completed haploinsufficiency and triplosensitivity review in ClinGen (accessed January 20, 2021, Supplemental Table 3). There are currently 38 regions that have sufficient evidence for haploinsufficiency and 21 regions with sufficient evidence for triplosensitivity. The majority of these regions contain at least one gene with a high NeuroSCORE (30/38 and 14/21, respectively). Next, we explored the NeuroSCORE profile of several recurrent CNV regions to help identify genes that may contribute to CNS-related phenotypic features.

The 7q11.23 recurrent region is approximately 1.5 megabases with deletions associated with Williams-Beuren syndrome (WBS) and gains associated with 7q11.23 duplication syndrome. The typical WBS deletion/duplication region contains 25 total genes, of which 15 are scored and six are high scoring genes. The *GTF2I* and *GTF2IRD1* genes have been implicated as key genes driving the neurobehavioral phenotype^59,60^, though *GTF2I* and *GTF2IRD1* knockout mice suggests that neither gene fully recapitulates the neurobehavioral aspects of WBS^59^. Aside from *GTF2I*, NeuroSCORE identified five high scoring genes (Figure 4), all with evidence for CNS involvement: *STX1A* has been associated with ASD^61^ and syndromic ID^62^ in humans, *LIMK1* sequence variants are associated with ASD^18^ and visuospatial impairment^63^, while *LIMK1* deficient mice have fewer cortical pyramidal neurons^64^, impaired long-term memory^65^ and spatial learning^66^; *BAZ1B* knockout mice show aberrant neurogenesis^67^ while clinical studies show variants associated with ASD^18^, Klippel-Feil syndrome^68^, and congenital heart defects^18^; *EIF4H* deficient mice have a smaller body size, behavioral impairments, and reduced brain volume^69^; finally, a single *CLIP2* variant has been associated with ASD^70^.

The 22q11.2 region is also associated with recurrent deletion and duplication syndromes. Within the typical 22q11.2 region there are 64 genes, of which 38 are scored and eight are high scoring genes. However, the 22q11.2 deletion/duplication syndrome region presents a challenge when interpreting CNVs that are smaller than the common breakpoint in the A-F deletion (breakpoints refer to areas of repetitive DNA that cause recurrent CNVs and often are given letter designations). Our analysis found the 22q11.2 A-B, B-D, and D-F breakpoint CNVs each harbor multiple scored genes and at least one high scoring gene (Figure 4), suggesting that CNVs of any of these smaller regions of 22q11.2 may be pathogenic for CNS-related clinical features. Two previous studies in cohorts of individuals with 22q11.2 deletion syndrome are also consistent with this finding. The first study analyzed neuroimaging data and transcriptomics and identified the *MAPK1* gene (a high scoring gene in the D-F region) as a potential driver gene of brain morphology changes^71^. The second study analyzed 22q11.2 deletion size and IQ score, finding that IQ score was partially explained by deletion size (A-B vs. B-D)^72^. Taken together, NeuroSCORE identified several candidate genes in the 7q11.23 and 22q11.2 regions, some of which are supported by other studies as well as additional genes that may provide new insight into the role of these regions in neurological phenotypes.

Emerging CNV syndromes also pose challenges for clinical interpretation. Applying NeuroSCORE to rare CNVs can help determine if they score in the same range as known pathogenic CNVs while also identifying candidate genes. We queried the ClinGen Dosage Sensitivity Map and identified the 2p16.1-p15 CNV as one with limited data on whether it caused a duplication syndrome. To date, seven individuals have been reported with a duplication and neurological clinical features^73–75^. Experimental studies in zebrafish of the previous candidate genes have implicated the *BCL11A, USP34, REL*, and *XPO1* genes^76^. Using NeuroSCORE, we first find this CNV has a median score of 30%, well above the average median score in our affected case gain CNVs of 16%. Furthermore, within this region are four high scoring (*BCL11A, USP34, XPO1*, and *CCT4*), of which the *CCT4* gene was not previously identified. Experimental work in *Drosophila* shows *CCT4* knockdown results in severely reduced dendritic growth^78^, as well as abnormalities of the eye and other organs^77^; furthermore, two *de novo*, missense variants have been reported in two individuals with ASD^18^. These data support a role for 2p15p16.1 gains in CNS phenotypes and indicate the *CCT4* gene as a new candidate gene.

## DISCUSSION

We hypothesized that CNS-related disease genes are likely to be identified by multiple metrics that assess different genic properties. After combining multiple genome-wide databases that assess different properties, we found that five properties were independently and positively associated with genes already known to cause CNS-related conditions as identified in the OMIM database. A total of 8,296 genes were identified by at least one of five gene metrics and 1,580 genes were identified by three or more. These high scoring genes are enriched in neurological processes in humans, developmental and neurological pathways, affected by CNVs more often in a neurodevelopmentally-affected cohort, and show neurologically related phenotypes in murine experiments (Table 3 and Figure 3). Of these 1,580 high scoring genes, 1,062 (67%) do not yet have any phenotype association in OMIM as of December 31, 2020 and 252 genes (15.9%) have no associated variants in the Human Genome Mutation Database (HGMD; accessed December 31, 2020). These 1,580 high scoring genes likely represent a significant proportion of the approximately 1,000 undiscovered neurodevelopmental disease genes proposed by a recent analysis^9^.

The findings from our pathway and GO analyses found significant enrichment in multiple neurological processes, many with known disease or phenotype associations (Supplemental Tables 1 and 2): the BAF (or SWI/SNF) complex^42^, the NuRD complex^43^, neuronal organization with microtubule tracking^44^, tau protein activity^45^, histone binding processes^46^, the Cajal body^47–49^, proteasome activity^50^, mRNA processing via both the spliceosome components^51,52^ and mRNA trafficking and binding^53,54^. Of these processes, splicing may be one of the most important. Tissue-specific splicing in the brain has shown high rates of alternative transcript splicing, suggesting that splicing proteins and proper splicing are imperative to neuronal development, structure, and function and appear to be evolutionarily conserved^55,56^. The Cajal body presents an interesting confluence of multiple previously discussed CNS-disease related processes as these nuclear bodies appear in fetal and neural cells to help mediate splicing, create parts of the spliceosome and ribonuclear proteins, activate transcription, and aid in chromatin and genome organization^47^. Considering that ASD and other neurodevelopmental disorders appear to begin in the fetal period^57,58^, the enrichment of Cajal body-associated genes raises an interesting target for additional study of genes related to possible “Cajalopathies”.

As of December 31^st^, 2020, our model identified 124 high scoring genes that are associated with conditions in OMIM that have no known CNS-related clinical features. While some of these genes will not be found to cause CNS-related features and are likely false positives, some have emerging evidence for causing CNS-related phenotypes. One example is *MORC2*, currently associated with only a form of Charcot-Marie-Tooth disease (OMIM #616661) but recently reported to cause a neurodevelopmental disorder^79^. Another example is *ANK2* (OMIM #106410), currently associated with only long QT syndrome type 4 and ankyrin-B-related cardiac arrhythmia (OMIM # 600919) though has variants associated with ASD^80^, ID^81^, and schizophrenia^82^. Using NeuroSCORE could help identify CNS disease genes that have been overlooked due to prior disease associations (see Supplementary Table 4 for a list of genes).

Finally, our model may be helpful to identify genes that influence or increase the risk for spectrum conditions, such as ASD. Multiple damaging variants in multiple scored genes could explain the risk or presence of a condition like ASD in individuals without a known variant in high-risk genes or common pathogenic CNVs. Damaging variants in scored genes may also help explain variable expressivity and reduced penetrance observed in many CNS-related genetic conditions (e.g., “two-hit” models).

Like Amirah Brigham’s and John Snow’s use of mapping in the 19^th^ century cholera epidemic, we have correlated existing data and created a new map to aid in discovery of conditions that broadly effect humanity. Our NeuroSCORE map of the human genome identifies and prioritizes potential disease genes of the CNS, which we validated using case-control and mouse model organism data. NeuroSCORE can be used for: bioinformatic analysis pipelines, identification of candidate disease genes in individuals with neurological phenotypes, guidance of basic and clinical research, the development of genetic tests, and furthering research on treatments for these conditions as current or future medications may target specific proteins or pathways. Future directions of model development can include identifying interaction terms to improve model precision as well as adding new metrics such as transcriptome profiles from brain expressed genes. While there are genes that cause CNS-related conditions not identified by NeuroSCORE (e.g., *GABRG2*), our model represents a potentially significant step forward in research to improve ultimately diagnostics for individuals with genetic causes of neurological conditions.

### Limitations

One limitation to this study and model is that it analyzes only protein-coding genes and excludes disease mechanisms such as mitochondrial, epigenetic, and disruption of enhancer and untranslated regions. Recent work in a small ASD cohort indicates that ASD risk may be influenced by variants in non-coding regions^83^. Similarly, genes causing autosomal recessive conditions are not well represented due to the use of gnomAD’s loss-of-function data. However, a recent analysis in the Deciphering Developmental Disorders cohort found approximately 3.6% of individuals from non-consanguineous families had a condition consistent with a recessive inheritance pattern^84^. Another limitation is that our outcome variable (CNS clinical features) is drawn from OMIM, which is an imperfect database of genotype-phenotype information with possibly inaccurate or outdated information and ascertainment bias. Finally, many conditions are not yet fully phenotyped, with rare phenotypes or age-related phenotypes not well represented.

## ACKNOWLEDGEMENTS

We would like to thank Dr. Lori Erby and Colleen Caleshu for helpful comments and thoughts on this project, as well as Dr. Aric Schadler for support with statistical methodology.

