## Supplemental Tables 1 - 4 for "NeuroSCORE: A Genome-wide Omics-Based Model to Identify Candidate Disease Genes of the Central Nervous System"

**Supplemental Table 1: Top Five Enriched Gene Ontology Terms & Candidate OMIM Genes Among High Scoring Genes**

| GO Annotation Term | Genes (N) | Genes Expected (N) | Enrichment | Genes without OMIM CNS-Related Phenotypes |
|---|---|---|---|---|
| **Biological Processes** | | | | |
| Positive regulation of protein localization to Cajal body (GO:1904871) | 9 | 0.7 | 13.0* | ***CCT2, CCT3, CCT4, CCT7, CCT8, CCT6A, DKC1, TCP1*** |
| Positive regulation of establishment of protein localization to telomere (GO:1904851) | 9 | 0.8 | 11.7* | ***CCT2, CCT3, CCT4, CCT7, CCT8, CCT6A, DKC1, TCP1*** |
| Axo-dendritic protein transport (GO:0099640) | 10 | 1.0 | 10.0* | *DLG2, KIF5B, MAP1A, TERF2,* ***SFPQ*** |
| Alternative mRNA splicing, via spliceosome (GO:0000380) | 13 | 1.3 | 9.9*** | ***CELF4***, ***DDX5***, *DDX17, DHX9, HNRNPM, RBM17,* ***SFPQ,*** *SFSWAP,* ***SLU7,*** *SRSF1* |
| Positive regulation of telomerase RNA localization to Cajal body (GO:1904874) | 11 | 1.2 | 9.5** | ***CCT2, CCT3, CCT4***, ***CCT7, CCT8, CCT6A, DKC1,*** *RUVBL1, RUVBL2,* ***TCP1*** |
| **Cellular Component** | | | | |
| nBAF complex (GO:0071565) | 12 | 1.2 | 10.4*** | *DPF1, SMARCC1* |
| Proteasome regulatory particle, base subcomplex (GO:0008540) | 9 | 0.9 | 9.8** | ***PSMC1, PSMC2, PSMC3, PSMC4, PSMC5, PSMC6, PSMD1, PSMD2, PSMD4*** |
| Chaperonin-containing T-complex (GO:0005832) | 8 | 0.9 | 9.5* | ***CCT2, CCT3, CCT4, CCT7, CCT8, CCT6A, TCP1*** |
| NuRD complex (GO:0016581) | 11 | 1.2 | 8.9** | *MTA3, HDAC2, RBBP7, MBD3, ZBTB7A, MTA1, CHD5, RBBP4* |
| Proteasome regulatory particle (GO:0005838) | 15 | 1.7 | 8.9*** | ***PSMC4, PSMC6***, *PSMD6, PSMD14,* ***PSMD2***, *PSMD11,* ***PSMC3***, ***PSMC1***, *PSMD13,* ***PSMC2***, *ADRM1,* ***PSMD1***, *PSMD3,* ***PSMD4, PSMC5*** |
| **Molecular Function** | | | | |
| Protein kinase A catalytic subunit binding (GO:0034236) | 10 | 1.0 | 10.0** | *CSK, EZR,* ***GSK3B, GSK3A***, *PRKAR1B, PRKAR2B, PJA2* |
| Tau-protein kinase activity (GO:0050321) | 14 | 1.7 | 8.3** | *TTBK1,* ***GSK3B***, *TAOK2,* ***GSK3A***, *BRSK1, MARK4, FYN, BRSK2, ROCK2, TAOK1, MARK2* |
| Microtubule plus-end binding (GO:0051010) | 10 | 1.2 | 8.1* | *CLASP1, CLASP2, CLIP1, CLIP2, MAPRE1, MAPRE2, NUMA1* |
| Pre-mRNA binding (GO:0036002) | 21 | 2.9 | 7.4*** | *CELF1, CELF2,* ***CELF4,*** *CELF5,* ***DDX5***, *HNRNPL, PRPF8, RBM22, RBM4, SF1, SRSF2,* ***SLU7***, *TRA2B, U2AF2* |
| Lysine-acetylated histone binding (GO:0070577) | 11 | 1.5 | 7.2* | *CARM1, BRD2, BRD3, BRD4, BRD7, ZMYND8* |

*$p \leq 0.05$, **$p \leq 0.01$, ***$p \leq 0.001$; *p*-values for Chi-squared testing uses Bonferroni correction for multiple testing for Biological processes (9,050 unique tests), Cellular Component (1,472 unique tests), Molecular Function (2,811 unique tests); **bold** genes appear two or more times in this table

**Supplemental Table 2: Select Pathways and Candidate Genes from AmiGO Visualization**

| GO Term | FDR | Genes without OMIM CNS-Related Phenotypes |
|---|---|---|
| Regulation of dendrite development (GO:0050773) | 4.1E$^{-23}$ | ***ACTR2,*** *ADGRB3*, ***ANAPC2, BAIAP2***, ***CAMK1D,*** *CAMSAP2*, ***CAPRIN1,*** *CARM1*, ***CRK, CYFIP1, DAB2IP, DBN1,*** *GSK3A*, *GSK3B*, ***PARP6, PREX1, STK11,*** *YWHAH* |
| Regulation of morphogenesis involved in differentiation (GO:0010769) | 8.0E$^{-34}$ | *ACTN4*, ***ACTR2***, ***ANAPC2,*** *ARHGEF7*, *ARPC2*, ***BAIAP2***, ***CAPRIN1***, *CORO1C*, ***CRK***, ***DBN1,*** *DMTN*, *P4HB*, ***PARP6, PREX1***, *PTK2*, ***PTPRD,*** *RCC2*, *TESK1* |
| Positive regulation of neurogenesis (GO:0050769) | 1.9E$^{-27}$ | ***ACTR2***, *AMIGO1*, ***ANAPC2, BAIAP2***, *BMPR2*, ***CAPRIN1, CYFIP1, DBN1***, *ETV5*, *FN1*, *HDAC2*, *HIF1A*, *LIMK1*, ***LRP1***, *NAP1L1*, ***NDEL1, NPTN***, *NUMBL*, ***PARP6***, *PLXNB1*, ***PLXNB2***, *PPP1CC*, *PRMT5*, ***PRPF19, PTPRD,*** *PTPRZ1*, *RAB11A*, *RELA*, *RHEB*, ***SNW1***, *SPEN*, *SRF*, ***STK11***, *TTBK1*, ***TWF2***, *XRCC5* |
| Positive regulation of neuron projection development (GO:0010976) | 1.8E$^{-22}$ | ***ACTR2,*** *AP2A1*, *APBB1*, *ARHGAP35*, ***BAIAP2, CAMK1D***, ***CAPRIN1, CYFIP1***, ***DAB2IP, DBN1,*** *DPYSL3*, *FYN*, ***LRP1,*** *MARK2*, ***NDEL1,*** *NDRG4*, ***NPTN***, ***PLXNB2***, *PPP2R5B*, *PTK2B*, *RAP1A*, *RAPGEF1*, ***SF3A2***, *STMN2*, ***TWF2*** |
| Regulation of mRNA stability (GO:0043488) | 5.8E$^{-30}$ | *ANP32A*, ***CELF1***, *DHX9*, *EIF4ENIF1*, *ELAVL1*, *FASTK*, ***FXR1***, ***FXR2,*** *HNRNPA0*, ***HNRNPC***, *HNRNPD*, ***HNRNPM***, ***HNRNPR***, ***HSPA8***, *KHSRP*, *LARP1*, *MAPKAPK2*, *NPM1*, ***PABPC1***, *PABPC4*, *PAIP1*, *PSMA1*, *PSMA2*, *PSMA3*, *PSMA5*, *PSMA6*, *PSMA7*, *PSMB1*, *PSMB3*, *PSMB5*, *PSMC1*, *PSMC2*, *PSMC3*, *PSMC4*, *PSMC5*, *PSMC6*, *PSMD1*, *PSMD11*, *PSMD13*, *PSMD14*, *PSMD2*, *PSMD3*, *PSMD4*, *PSMD6*, *PSME3*, *PUM2*, *ROCK2*, *SAMD4B*, *SERBP1*, ***SYNCRIP***, ***THRAP3***, *UBC*, *UPF1*, *XPO1*, *YTHDF1*, *YTHDF2*, *YTHDF3*, *YWHAB*, *YWHAZ* |
| Regulation of mRNA splicing via the spliceosome (GO:0048024) | 7.6E$^{-36}$ | ***CELF1***, *CELF2*, *CELF4*, *CELF5*, *DAZAP1*, *DDX17*, ***DDX5***, ***FXR1***, ***FXR2,*** *HNRNPL*, ***HSPA8***, *IK*, *KHDRBS3*, ***MAGOH***, *NCBP1*, *NCL*, *NOVA1*, ***PRPF19***, *QKI*, *RBFOX1*, *RBFOX2*, *RBM17*, *RBM25*, *RBM39*, *RBM4*, *RBM5*, *RNPS1*, *SART3*, *SF1*, *SF3B4*, *SFSWAP*, *SMU1*, *SNRNP70*, ***SNW1***, *SRPK2*, ***SRSF1***, *SRSF2*, *SRSF3*, *SRSF4*, *SRSF7*, *SRSF9*, ***THRAP3***, *TRA2B*, *U2AF2*, *WTAP*, *YTHDC1*, *ZBTB7A* |
| Catalytic step two of the spliceosome (GO:0071013) | 1.3E$^{-22}$ | *CDC5L*, *CWC15*, *DDX23*, ***DDX5,*** *HNRNPA3*, ***HNRNPC***, *HNRNPF*, *HNRNPH1*, ***HNRNPM***, ***HNRNPR, MAGOH***, ***PABPC1,*** *PLRG1*, ***PRPF19***, *PRPF4B*, *PRPF6*, *PRPF8*, *RBM22*, *SART1*, *SF3A1*, ***SF3A2***, *SF3A3*, *SF3B1*, *SF3B2*, *SF3B3*, *SLU7*, *SNRNP200,SNRPD3*, ***SNW1,*** *SRRM1*, *SRRM2*, ***SRSF1***, ***SYNCRIP***, *XAB2* |

FDR: false discovery rate; **bold** genes are represented two or more times in this table.

**Pathway Analysis of High Scoring Genes via Cytoscape Visualization**

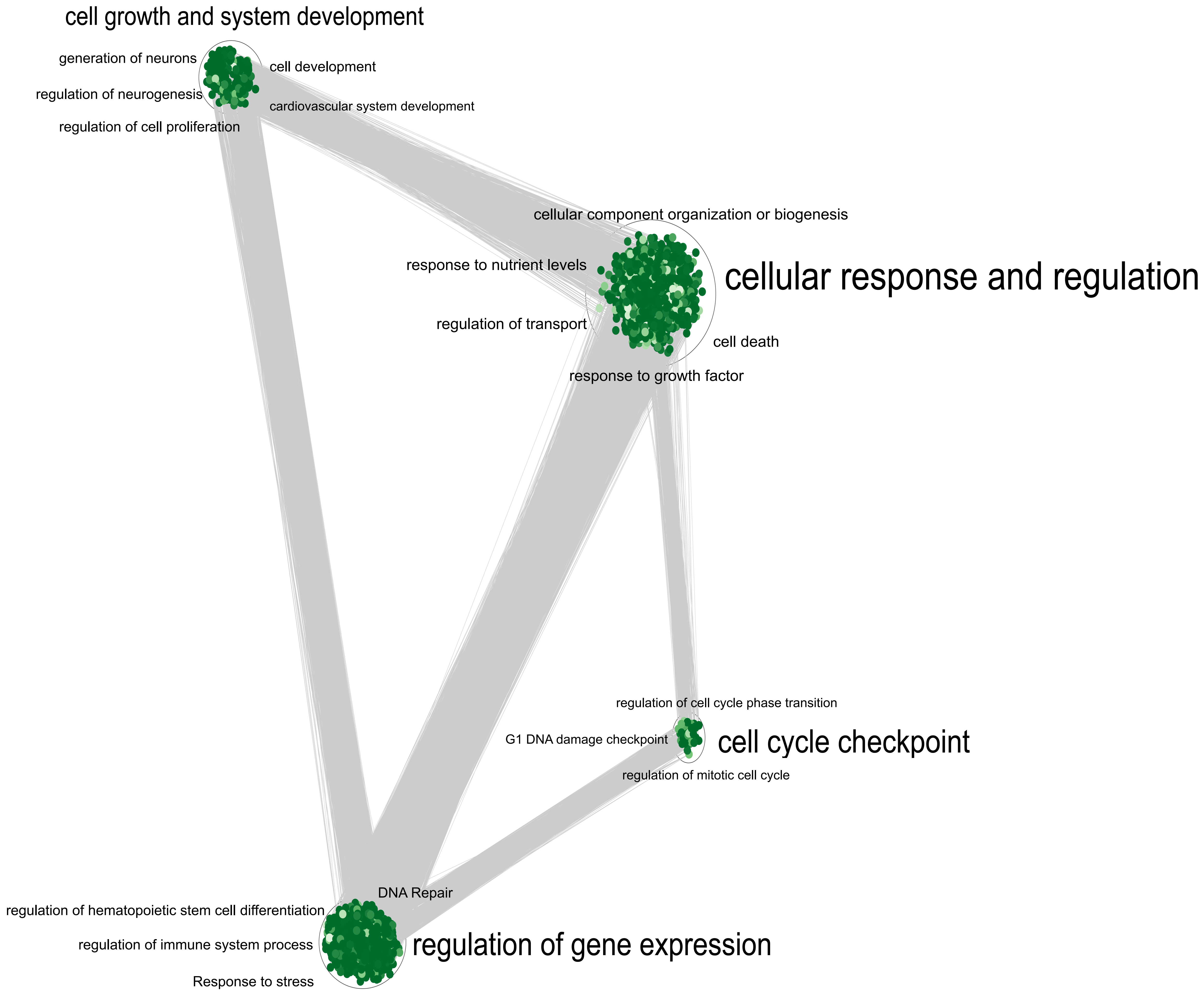


**Node color represents the *p*-value with darker shades corresponding to lower *p*-values; node size represents odds ratio with increasing node size illustrating increasing odds.**

Supplementary Table 3: ClinGen Dosage Sensitivity with NeruoSCORE by Gene

| ClinGen Region | | ISCA ID | Haploinsufficiency | Triplosensitivity | Coordinates (GRCh37/hg19) |
|---|---|---|---|---|---|
| 1p36 terminal region (includes GABRD) | | ISCA-37434 | 3 | 2 | chr1:834,083-6,289,973 |
| **Gene** | **NeuroSCORE** | **Gene** | **NeuroSCORE** | **Gene** | **NeuroSCORE** |
| *SAMD11* | 0 | *MRPL20* | 2 | *RER1* | 2 |
| *AL645608.1* | NA | *ANKRD65* | 1 | *PEX10* | 2 |
| *NOC2L* | 2 | *TMEM88B* | 1 | *PLCH2* | 1 |
| *KLHL17* | 0 | *VWA1* | 1 | *PANK4* | 1 |
| *PLEKHN1* | 0 | *ATAD3C* | 0 | *HES5* | 0 |
| *PERM1* | 0 | *ATAD3B* | 1 | *TNFRSF14* | 0 |
| *HES4* | 1 | *ATAD3A* | 0 | *FAM213B* | 2 |
| *ISG15* | 1 | *TMEM240* | 2 | *MMEL1* | 1 |
| *AGRN* | 2 | *SSU72* | 3 | *TTC34* | 0 |
| *RNF223* | 0 | *AL645728.1* | 0 | *ACTRT2* | 0 |
| *C1orf159* | 0 | *FNDC10* | 1 | *PRDM16* | 2 |
| *TTLL10* | 0 | *MIB2* | 1 | *ARHGEF16* | 0 |
| *TNFRSF18* | 0 | *MMP23B* | 0 | *MEGF6* | 0 |
| *TNFRSF4* | 0 | *CDK11B* | 2 | *TPRG1L* | 2 |
| *SDF4* | 2 | *SLC35E2B* | 1 | *WRAP73* | 1 |
| *B3GALT6* | 1 | *CDK11A* | 0 | *TP73* | 1 |
| *C1QTNF12* | 0 | *SLC35E2* | 0 | *CCDC27* | 0 |
| *UBE2J2* | 2 | *NADK* | 2 | *SMIM1* | 0 |
| *SCNN1D* | 1 | *GNB1* | 3 | *LRRC47* | 2 |
| *ACAP3* | 2 | *CALML6* | 0 | *CEP104* | 0 |
| *PUSL1* | 0 | *TMEM52* | 0 | *DFFB* | 0 |
| *INTS11* | 2 | *CFAP74* | 0 | *C1orf174* | 0 |
| *CPTP* | 2 | *GABRD* | 2 | *AJAP1* | 1 |
| *TAS1R3* | 0 | *PRKCZ* | 2 | *NPHP4* | 1 |
| *DVL1* | 2 | *FAAP20* | 1 | *KCNAB2* | 1 |
| *MXRA8* | 2 | *AL590822.2* | NA | *AL035406.1* | NA |
| *AURKAIP1* | 2 | *AL590822.1* | NA | *CHD5* | 3 |
| *CCNL2* | 1 | *SKI* | 3 | *RPL22* | 2 |
| *RP4-758J18.2* | 1 | *MORN1* | 0 | *RNF207* | 0 |
| | | | | *ICMT* | 1 |

Supplementary Table 3: ClinGen Dosage Sensitivity with NeruoSCORE by Gene

| ClinGen Region | ISCA ID | Haploinsufficiency | Triplosensitivity | Coordinates (GRCh37/hg19) |
|---|---|---|---|---|
| 1q21.1 recurrent (TAR syndrome) region (BP2-BP3, proximal) (includes RBM8A) | ISCA-37428 | 1 | 1 | chr1:145,386,507-145,748,064 |

| Gene | NeuroSCORE | Gene | NeuroSCORE |
|---|---|---|---|
| *HFE2* | 0 | *ANKRD35* | 0 |
| *TXNIP* | 2 | *PIAS3* | 2 |
| *POLR3GL* | 2 | *NUDT17* | 0 |
| *ANKRD34A* | 2 | *POLR3C* | 0 |
| *LIX1L* | 1 | *RNF115* | 0 |
| *RBM8A* | 2 | *CD160* | 0 |
| *PEX11B* | 2 | *PDZK1* | 0 |
| *ITGA10* | 0 | | |

| ClinGen Region | ISCA ID | Haploinsufficiency | Triplosensitivity | Coordinates (GRCh37/hg19) |
|---|---|---|---|---|
| 1q21.1 recurrent region (BP3-BP4, distal) (includes GJA5) | ISCA-37421 | 3 | 3 | chr1:146,577,486-147,394,506 |

| Gene | NeuroSCORE |
|---|---|
| *PRKAB2* | 0 |
| *FMO5* | 0 |
| *CHD1L* | 2 |
| *BCL9* | 0 |
| *ACP6* | 0 |
| *GJA5* | 0 |
| *GJA8* | 0 |

Supplementary Table 3: ClinGen Dosage Sensitivity with NeruoSCORE by Gene

| ClinGen Region | | ISCA ID | Haploinsufficiency | Triplosensitivity | Coordinates (GRCh37/hg19) |
|---|---|---|---|---|---|
| 1q21.2 region (polymorphic region)(non-unique; also maps to chromosomes 16, 7, and others in GRCh38) | | ISCA-37469 | Haploinsufficiency unlikely | Triplosensitivity unlikely | chr1:148,867,551-149,768,855 |
| **Gene** | **NeuroSCORE** | | | | |
| *HIST2H3PS2* | 0 | | | | |
| *FAM72C* | 0 | | | | |
| *FCGR1A* | 0 | | | | |
| *HIST2H2BF* | 0 | | | | |

| ClinGen Region | ISCA ID | Haploinsufficiency | Triplosensitivity | Coordinates (GRCh37/hg19) |
|---|---|---|---|---|
| 1q43q44 terminal region (includes AKT3) | ISCA-37493 | 3 | 0 | chr1:243,287,730-245,318,287 |
| **Gene** | **NeuroSCORE** | **Gene** | **NeuroSCORE** | |
| *CEP170* | 1 | *CATSPERE* | 0 | |
| *SDCCAG8* | 1 | *DESI2* | 0 | |
| *AKT3* | 4 | *COX20* | 1 | |
| *ZBTB18* | 5 | *HNRNPU* | 5 | |
| *C1orf100* | 0 | *RP11-156E8.1* | 0 | |
| *ADSS2* | 2 | *EFCAB2* | 0 | |

| ClinGen Region | ISCA ID | Haploinsufficiency | Triplosensitivity | Coordinates (GRCh37/hg19) |
|---|---|---|---|---|
| 2p24.3 MYCN-DDX1 duplication region | ISCA-46287 | Not Yet Evaluated | 2 | chr2:15,708,677-16,185,337 |
| **Gene** | **NeuroSCORE** | | | |
| *DDX1* | 4 | | | |
| *AC008271.1* | 0 | | | |
| *MYCN* | 0 | | | |

Supplementary Table 3: ClinGen Dosage Sensitivity with NeruoSCORE by Gene

| ClinGen Region | ISCA ID | Haploinsufficiency | Triplosensitivity | Coordinates (GRCh37/hg19) |
|---|---|---|---|---|
| 2p21 region (includes PREPL and SLC3A1) | ISCA-37440 | Gene associated with an AR phenotype | 0 | chr2:44,410,272-44,589,641 |
| **Gene** | **NeuroSCORE** | | | |
| *PPM1B* | 2 | | | |
| *SLC3A1* | 1 | | | |
| *PREPL* | 2 | | | |
| *CAMKMT* | 0 | | | |

| ClinGen Region | ISCA ID | Haploinsufficiency | Triplosensitivity | Coordinates (GRCh37/hg19) |
|---|---|---|---|---|
| 2p15p16.1 region (includes BCL11A) | ISCA-37408 | 3 | 1 | chr2:59,139,200-62,488,871 |
| **Gene** | **NeuroSCORE** | **Gene** | **NeuroSCORE** | |
| *BCL11A* | 3 | *AHSA2* | 1 | |
| *PAPOLG* | 2 | *USP34* | 3 | |
| *REL* | 1 | *XPO1* | 4 | |
| *PUS10* | 0 | *FAM161A* | 0 | |
| *PEX13* | 0 | *CCT4* | 3 | |
| *KIAA1841* | 0 | *COMMD1* | 2 | |
| *C2orf74* | 1 | *B3GNT2* | 2 | |

Supplementary Table 3: ClinGen Dosage Sensitivity with NeruoSCORE by Gene

| ClinGen Region | | ISCA ID | Haploinsufficiency | Triplosensitivity | Coordinates (GRCh37/hg19) |
|---|---|---|---|---|---|
| 2q11.2 recurrent region (includes ARID5A, LMAN2L) | | ISCA-37495 | 1 | 1 | chr2:96,739,012-97,671,429 |
| **Gene** | **NeuroSCORE** | **Gene** | **NeuroSCORE** | | |
| *ADRA2B* | 0 | *ARID5A* | 0 | | |
| *ASTL* | 0 | *KANSL3* | 3 | | |
| *DUSP2* | 0 | *LMAN2L* | 1 | | |
| *STARD7* | 2 | *CNNM4* | 0 | | |
| *TMEM127* | 2 | *CNNM3* | 0 | | |
| *CIAO1* | 2 | *ANKRD23* | 1 | | |
| *SNRNP200* | 4 | *ANKRD39* | 1 | | |
| *ITPRIPL1* | 0 | *SEMA4C* | 2 | | |
| *NCAPH* | 0 | *FAM178B* | 0 | | |

| ClinGen Region | ISCA ID | Haploinsufficiency | Triplosensitivity | Coordinates (GRCh37/hg19) |
|---|---|---|---|---|
| 2q13 recurrent region (includes NPHP1) | ISCA-37405 | Gene associated with AR phenotype | Triplosensitivity unlikely | chr2:110,862,108-110,983,703 |
| **Gene** | **NeuroSCORE** | | | |
| *MALL* | 0 | | | |
| *NPHP1* | 0 | | | |

| ClinGen Region | ISCA ID | Haploinsufficiency | Triplosensitivity | Coordinates (GRCh37/hg19) |
|---|---|---|---|---|
| 2q13 recurrent region (includes BCL2L11) | ISCA-37496 | 2 | 2 | chr2:111,392,193-113,104,742 |
| **Gene** | **NeuroSCORE** | **Gene** | **NeuroSCORE** | |
| *BUB1* | 0 | *MERTK* | 0 | |
| *ACOXL* | 0 | *TMEM87B* | 0 | |
| *BCL2L11* | 0 | *FBLN7* | 0 | |
| *ANAPC1* | 1 | *ZC3H6* | 1 | |

Supplementary Table 3: ClinGen Dosage Sensitivity with NeruoSCORE by Gene

| ClinGen Region | | ISCA ID | Haploinsufficiency | Triplosensitivity | Coordinates (GRCh37/hg19) |
|---|---|---|---|---|---|
| 2q21.1 recurrent region (includes ARHGEF4, GPR148) | | ISCA-46288 | 1 | 0 | chr2:131,477,509-131,929,693 |
| **Gene** | **NeuroSCORE** | **Gene** | **NeuroSCORE** | | |
| *GPR148* | 0 | *FAM168B* | 2 | | |
| *AMER3* | 0 | *PLEKHB2* | 3 | | |
| *ARHGEF4* | 2 | | | | |

| ClinGen Region | | ISCA ID | Haploinsufficiency | Triplosensitivity | Coordinates (GRCh37/hg19) |
|---|---|---|---|---|---|
| 2q37.3 terminal region (includes HDAC4) | | ISCA-37394 | 3 | 0 | chr2:239,954,693-242,930,600 |
| **Gene** | **NeuroSCORE** | **Gene** | **NeuroSCORE** | **Gene** | **NeuroSCORE** |
| *HDAC4* | 2 | *GPR35* | 0 | *THAP4* | 3 |
| *AC017028.1* | NA | *AQP12B* | 0 | *ATG4B* | 4 |
| *AC062017.1* | 0 | *AQP12A* | 0 | *DTYMK* | 0 |
| *AC079612.1* | 0 | *KIF1A* | 5 | *ING5* | 1 |
| *AC093802.1* | 0 | *AGXT* | 0 | *D2HGDH* | 0 |
| *NDUFA10* | 2 | *C2orf54* | 0 | *GAL3ST2* | 0 |
| *OR6B2* | 0 | *AC104809.3* | 0 | *NEU4* | 0 |
| *PRR21* | 0 | *SNED1* | 1 | *PDCD1* | 0 |
| *OR6B3* | 0 | *MTERF4* | 0 | *CXXC11* | 1 |
| *COPS9* | 2 | *PASK* | 0 | *AC131097.4* | 0 |
| *OTOS* | 0 | *PPP1R7* | 3 | | |
| *GPC1* | 2 | *ANO7* | 0 | | |
| *AC110619.2* | 0 | *HDLBP* | 3 | | |
| *ANKMY1* | 0 | *SEPTIN2* | 1 | | |
| *DUSP28* | 0 | *FARP2* | 0 | | |
| *RNPEPL1* | 0 | *STK25* | 2 | | |
| *CAPN10* | 1 | *BOK* | 2 | | |

Supplementary Table 3: ClinGen Dosage Sensitivity with NeruoSCORE by Gene

| ClinGen Region | ISCA ID | Haploinsufficiency | Triplosensitivity | Coordinates (GRCh37/hg19) |
|---|---|---|---|---|
| 1 copy: 2q telomere\|3 copies: 2q telomere | ISCA-37470 | Haploinsufficiency unlikely | Triplosensitivity unlikely | chr2:242,930,600-243,102,476 |

| ClinGen Region | ISCA ID | Haploinsufficiency | Triplosensitivity | Coordinates (GRCh37/hg19) |
|---|---|---|---|---|
| 3q29 recurrent region (includes DLG1) | ISCA-37443 | 3 | 2 | chr3:195,756,054-197,344,662 |

| Gene | NeuroSCORE | Gene | NeuroSCORE |
|---|---|---|---|
| *TFRC* | 2 | *NRROS* | 0 |
| *ZDHHC19* | 0 | *PIGX* | 2 |
| *SLC51A* | 0 | *CEP19* | 0 |
| *PCYT1A* | 1 | *PAK2* | 3 |
| *TCTEX1D2* | 2 | *SENP5* | 3 |
| *TM4SF19* | 1 | *NCBP2* | 2 |
| *UBXN7* | 2 | *PIGZ* | 1 |
| *RNF168* | 0 | *MELTF* | 0 |
| *SMCO1* | 0 | *DLG1* | 3 |
| *WDR53* | 0 | *BDH1* | 1 |
| *FBXO45* | 2 | | |

Supplementary Table 3: ClinGen Dosage Sensitivity with NeruoSCORE by Gene

| ClinGen Region | ISCA ID | Haploinsufficiency | Triplosensitivity | Coordinates (GRCh37/hg19) |
|---|---|---|---|---|
| 4p16.3 terminal (Wolf-Hirshhorn syndrome) | ISCA-37429 | 3 | 2 | chr4:331,568-2,010,962 |

| Gene | NeuroSCORE | Gene | NeuroSCORE |
|---|---|---|---|
| *ZNF141* | 0 | *RNF212* | 0 |
| *ZNF721* | 0 | *SPON2* | 0 |
| *PIGG* | 0 | *CTBP1* | 4 |
| *PDE6B* | 1 | *MAEA* | 2 |
| *ATP5I* | 2 | *UVSSA* | 0 |
| *MYL5* | 0 | *CRIPAK* | 0 |
| *MFSD7* | 0 | *NKX1-1* | 0 |
| *PCGF3* | 3 | *FAM53A* | 0 |
| *CPLX1* | 1 | *SLBP* | 2 |
| *GAK* | 2 | *TMEM129* | 1 |
| *TMEM175* | 1 | *TACC3* | 0 |
| *DGKQ* | 1 | *FGFR3* | 2 |
| *SLC26A1* | 0 | *LETM1* | 2 |
| *IDUA* | 0 | *NSD2* | 4 |
| *FGFRL1* | 1 | *NELFA* | 0 |

Supplementary Table 3: ClinGen Dosage Sensitivity with NeruoSCORE by Gene

| ClinGen Region | ISCA ID | Haploinsufficiency | Triplosensitivity | Coordinates (GRCh37/hg19) |
|---|---|---|---|---|
| 5p15 terminal (Cri du chat syndrome) | ISCA-37390 | 3 | 2 | chr5:37,693-11,347,262 |

| Gene | NeuroSCORE | Gene | NeuroSCORE |
|---|---|---|---|
| *PLEKHG4B* | 0 | *IRX4* | 1 |
| *LRRC14B* | 0 | *IRX2* | 0 |
| *CCDC127* | 0 | *C5orf38* | 0 |
| *SDHA* | 2 | *IRX1* | 0 |
| *PDCD6* | 2 | *ADAMTS16* | 0 |
| *AHRR* | 1 | *ICE1* | 1 |
| *C5orf55* | 0 | *MED10* | 2 |
| *EXOC3* | 3 | *UBE2QL1* | 2 |
| *CTD-2228K2.5* | 0 | *NSUN2* | 1 |
| *SLC9A3* | 1 | *SRD5A1* | 0 |
| *CEP72* | 0 | *PAPD7* | 2 |
| *TPPP* | 1 | *ADCY2* | 3 |
| *ZDHHC11* | 0 | *C5orf49* | 0 |
| *ZDHHC11B* | 1 | *MTRR* | 0 |
| *BRD9* | 2 | *FASTKD3* | 0 |
| *TRIP13* | 1 | *SEMA5A* | 0 |
| *RP11-661C8.3* | 0 | *TAS2R1* | 0 |
| *NKD2* | 0 | *ATPSCKMT* | 0 |
| *SLC12A7* | 0 | *CCT5* | 3 |
| *SLC6A19* | 0 | *CMBL* | 1 |
| *SLC6A18* | 0 | *MARCHF6* | 3 |
| *TERT* | 2 | *ROPN1L* | 0 |
| *CLPTM1L* | 2 | *RP11-1C1.5* | 0 |
| *SLC6A3* | 1 | *ANKRD33B* | 0 |
| *LPCAT1* | 2 | *DAP* | 2 |
| *MRPL36* | 2 | *CTNND2* | 3 |
| *NDUF36* | 2 | | |

Supplementary Table 3: ClinGen Dosage Sensitivity with NeruoSCORE by Gene

| ClinGen Region | | ISCA ID | Haploinsufficiency | Triplosensitivity | Coordinates (GRCh37/hg19) |
|---|---|---|---|---|---|
| 5q35 recurrent (Sotos syndrome) region (includes NSD1) | | ISCA-37425 | 3 | 3 | chr5:175,728,979-177,047,793 |
| **Gene** | **NeuroSCORE** | **Gene** | **NeuroSCORE** | | |
| *SIMC1* | 0 | *NSD1* | 3 | | |
| *KIAA1191* | 2 | *RAB24* | 2 | | |
| *ARL10* | 0 | *MXD3* | 1 | | |
| *NOP16* | 1 | *PRELID1* | 2 | | |
| *HIGD2A* | 2 | *LMAN2* | 2 | | |
| *CLTB* | 2 | *RGS14* | 1 | | |
| *FAF2* | 3 | *SLC34A1* | 0 | | |
| *RNF44* | 2 | *PFN3* | 0 | | |
| *CDHR2* | 1 | *F12* | 0 | | |
| *GPRIN1* | 2 | *GRK6* | 2 | | |
| *SNCB* | 2 | *PRR7* | 1 | | |
| *EIF4E1B* | 0 | *DBN1* | 3 | | |
| *TSPAN17* | 2 | *PDLIM7* | 1 | | |
| *UNC5A* | 2 | *DOK3* | 0 | | |
| *HK3* | 0 | *DDX41* | 2 | | |
| *UIMC1* | 0 | *FAM193B* | 2 | | |
| *ZNF346* | 0 | *TMED9* | 2 | | |
| *FGFR4* | 0 | *B4GALT7* | 1 | | |

| ClinGen Region | ISCA ID | Haploinsufficiency | Triplosensitivity | Coordinates (GRCh37/hg19) |
|---|---|---|---|---|
| 1 copy: 6p telomere \| 3 copies: 6p telomere | ISCA-37471 | Haploinsufficiency unlikely | Triplosensitivity unlikely | chr6:259,528-339,802 |
| **Gene** | **NeuroSCORE** | | | |
| *DUSP22* | 1 | | | |

Supplementary Table 3: ClinGen Dosage Sensitivity with NeruoSCORE by Gene

| ClinGen Region | ISCA ID | Haploinsufficiency | Triplosensitivity | Coordinates (GRCh37/hg19) |
|---|---|---|---|---|
| 6q24 region (includes PLAGL1) | ISCA-37442 | 1 | 3 | chr6:144,243,292-144,416,561 |
| **Gene** | **NeuroSCORE** | | | |
| *ZC2HC1B* | 0 | | | |
| *PLAGL1* | 1 | | | |
| *SF3B5* | 1 | | | |

| ClinGen Region | | ISCA ID | Haploinsufficiency | Triplosensitivity | Coordinates (GRCh37/hg19) |
|---|---|---|---|---|---|
| 7q11.23 recurrent (Williams-Beuren syndrome) region (includes ELN) | | ISCA-37392 | 3 | 3 | chr7:72,744,455-74,142,510 |
| **Gene** | **NeuroSCORE** | **Gene** | **NeuroSCORE** | | |
| *FKBP6* | 0 | *CLDN3* | 0 | | |
| *FZD9* | 0 | *CLDN4* | 0 | | |
| *BAZ1B* | 3 | *METTL27* | 0 | | |
| *BCL7B* | 2 | *TMEM270* | 0 | | |
| *TBL2* | 0 | *ELN* | 0 | | |
| *MLXIPL* | 1 | *LIMK1* | 4 | | |
| *VPS37D* | 1 | *EIF4H* | 3 | | |
| *DNAJC30* | 1 | *LAT2* | 0 | | |
| *BUD23* | 2 | *RFC2* | 1 | | |
| *STX1A* | 3 | *CLIP2* | 4 | | |
| *ABHD11* | 1 | *GTF2IRD1* | 1 | | |
| | | *GTF2I* | 3 | | |

Supplementary Table 3: ClinGen Dosage Sensitivity with NeruoSCORE by Gene

| ClinGen Region | | ISCA ID | Haploinsufficiency | Triplosensitivity | Coordinates (GRCh37/hg19) |
|---|---|---|---|---|---|
| 7q11.23 recurrent distal region (includes HIP1, YWHAG) | | ISCA-46291 | 2 | 1 | chr7:75,158,048-76,063,176 |
| **Gene** | **NeuroSCORE** | **Gene** | **NeuroSCORE** | | |
| *HIP1* | 2 | *STYXL1* | 1 | | |
| *CCL26* | 0 | *MDH2* | 2 | | |
| *CCL24* | 0 | *SRRM3* | 1 | | |
| *RHBDD2* | 2 | *HSPB1* | 2 | | |
| *POR* | 2 | *YWHAG* | 3 | | |
| *UQCRHL* | NA | *SRCRB4D* | 0 | | |
| *TMEM120A* | NA | *ZP3* | 0 | | |

| ClinGen Region | | ISCA ID | Haploinsufficiency | Triplosensitivity | Coordinates (GRCh37/hg19) |
|---|---|---|---|---|---|
| 7q36.3 ZRS (SHH cis-regulatory) duplication region (within LMBR1 intron 5) | | ISCA-37467 | 0 | 3 | chr7:156,583,796-156,584,568 |
| **Gene** | **NeuroSCORE** | | | | |
| *LMBR1* | 1 | | | | |

| ClinGen Region | ISCA ID | Haploinsufficiency | Triplosensitivity | Coordinates (GRCh37/hg19) |
|---|---|---|---|---|
| 8p23.1 region (DEFB gene cluster) | ISCA-37472 | Haploinsufficiency unlikely | Triplosensitivity unlikely | chr8:7,053,186-8,130,689 |
| **Gene** | **NeuroSCORE** | **Gene** | **NeuroSCORE** | |
| *ZNF705G* | 0 | *PRR23D1* | 0 | |
| *DEFB4B* | 0 | *DEFB107A* | 0 | |
| *SPAG11B* | 0 | *DEFB105A* | 0 | |
| *DEFB104B* | 0 | *DEFB106A* | 0 | |
| *DEFB106B* | 0 | *DEFB104A* | 0 | |
| *DEFB105B* | 0 | *SPAG11A* | 0 | |
| *DEFB107B* | 0 | *ZNF705B* | 0 | |

Supplementary Table 3: ClinGen Dosage Sensitivity with NeruoSCORE by Gene

| ClinGen Region | | ISCA ID | Haploinsufficiency | Triplosensitivity | Coordinates (GRCh37/hg19) |
|---|---|---|---|---|---|
| 10q22.3q23.2 recurrent region (LCR-3/4-flanked) (includes BMPR1A) | | ISCA-37424 | 3 | 1 | chr10:81,682,843-88,739,388 |
| **Gene** | **NeuroSCORE** | **Gene** | **NeuroSCORE** | **Gene** | **NeuroSCORE** |
| *SFTPD* | 0 | *TSPAN14* | 2 | *CCSER2* | 2 |
| *TMEM254* | 0 | *SH2D4B* | 0 | *GRID1* | 1 |
| *PLAC9* | 0 | *NRG3* | 0 | *WAPAL* | 2 |
| *ANXA11* | 1 | *GHITM* | 2 | *OPM4* | 0 |
| *AL359195.1* | NA | *C10orf99* | 0 | *LDB3* | 0 |
| *MAT1A* | 0 | *CDHR1* | 0 | *BMPR1A* | 1 |
| *DYDC1* | 0 | *LRIT2* | 0 | *MMRN2* | 0 |
| *DYDC2* | 0 | *LRIT1* | 0 | *SNCG* | 1 |
| *FAM213A* | 2 | *RGR* | 0 | *ADIRF* | 0 |

| ClinGen Region | | ISCA ID | Haploinsufficiency | Triplosensitivity | Coordinates (GRCh37/hg19) |
|---|---|---|---|---|---|
| 11p13 (WAGR syndrome) region | | ISCA-37401 | 3 | 1 | chr11:31,803,509-32,510,988 |
| **Gene** | **NeuroSCORE** | | | | |
| *ELP4* | 0 | | | | |
| *PAX6* | 3 | | | | |
| *RCN1* | 1 | | | | |
| *WT1* | 1 | | | | |

Supplementary Table 3: ClinGen Dosage Sensitivity with NeruoSCORE by Gene

| ClinGen Region | | ISCA ID | Haploinsufficiency | Triplosensitivity | Coordinates (GRCh37/hg19) |
|---|---|---|---|---|---|
| 11p11.2 (Potocki-Shaffer syndrome) region (includes ALX4, EXT2) | | ISCA-37441 | 3 | 0 | chr11:43,894,800-46,152,450 |
| **Gene** | **NeuroSCORE** | **Gene** | **NeuroSCORE** | **Gene** | **NeuroSCORE** |
| *ALKBH3* | 1 | *TSPAN18* | 0 | *CRY2* | 2 |
| *C11orf96* | 1 | *TP53I11* | 3 | *MAPK8IP1* | 3 |
| *ACCSL* | 0 | *PRDM11* | 1 | *C11orf94* | 0 |
| *ACCS* | 0 | *SYT13* | 2 | *PEX16* | 2 |
| *EXT2* | 2 | *CHST1* | 2 | *LARGE2* | 0 |
| *ALX4* | 0 | *CTD-2210P24.4* | 0 | *PHF21A* | 3 |
| *CD82* | 1 | *SLC35C1* | 0 | | |

| ClinGen Region | | ISCA ID | Haploinsufficiency | Triplosensitivity | Coordinates (GRCh37/hg19) |
|---|---|---|---|---|---|
| 11q13.2q13.4 recurrent region (includes SHANK2, FGFs) | | ISCA-37498 | 2 | 0 | chr11:67,763,646-71,236,931 |
| **Gene** | **NeuroSCORE** | **Gene** | **NeuroSCORE** | **Gene** | **NeuroSCORE** |
| *UNC93B1* | 0 | *MTL5* | 0 | *FGF19* | 0 |
| *ALDH3B1* | 0 | *CPT1A* | 1 | *FGF4* | 0 |
| *NDUFS8* | 2 | *MRPL21* | 2 | *FGF3* | 0 |
| *TCIRG1* | 0 | *IGHMBP2* | 0 | *ANO1* | 1 |
| *CHKA* | 2 | *MRGPRD* | 0 | *FADD* | 0 |
| *KMT5B* | 3 | *MRGPRF* | 0 | *PPFIA1* | 1 |
| *C11orf24* | 0 | *TPCN2* | 0 | *CTTN* | 2 |
| *LRP5* | 0 | *MYEOV* | 0 | *SHANK2* | 0 |
| *PPP6R3* | 3 | *CCND1* | 1 | *DHCR7* | 2 |
| *GAL* | 0 | *ORAOV1* | 0 | *NADSYN1* | 0 |

Supplementary Table 3: ClinGen Dosage Sensitivity with NeruoSCORE by Gene

| ClinGen Region | ISCA ID | Haploinsufficiency | Triplosensitivity | Coordinates (GRCh37/hg19) |
|---|---|---|---|---|
| 14q11.2 region including CHD8 and SUPT16H | ISCA-37438 | 2 | 0 | chr14:21,826,900-21,861,987 |

| Gene | NeuroSCORE |
|---|---|
| *SUPT16H* | 4 |
| *CHD8* | 5 |

| ClinGen Region | ISCA ID | Haploinsufficiency | Triplosensitivity | Coordinates (GRCh37/hg19) |
|---|---|---|---|---|
| 14q11.2 region (TCRA region) | ISCA-37476 | Haploinsufficiency unlikely | Triplosensitivity unlikely | chr14:22,111,109-23,021,097 |

| Gene | NeuroSCORE |
|---|---|
| *OR4E2* | 0 |

| ClinGen Region | ISCA ID | Haploinsufficiency | Triplosensitivity | Coordinates (GRCh37/hg19) |
|---|---|---|---|---|
| 14q32 region associated with UPD(14) phenotypes | ISCA-37449 | 2 | 0 | chr14:100,394,594-101,504,529 |

| Gene | NeuroSCORE | Gene | NeuroSCORE |
|---|---|---|---|
| *EML1* | 1 | *WARS* | 3 |
| *EVL* | 3 | *WDR25* | 0 |
| *DEGS2* | 0 | *BEGAIN* | 3 |
| *YY1* | 4 | *DLK1* | 0 |
| *SLC25A29* | 2 | *RTL1* | 1 |
| *SLC25A47* | 0 | | |

Supplementary Table 3: ClinGen Dosage Sensitivity with NeruoSCORE by Gene

| ClinGen Region | ISCA ID | Haploinsufficiency | Triplosensitivity | Coordinates (GRCh37/hg19) |
|---|---|---|---|---|
| DLK1-MEG3 Intergenic DMR | ISCA-37447 | 1 | 0 | chr14:101,191,391-101,294,616 |

| Gene | NeuroSCORE |
|---|---|
| *DLK1* | 0 |

| ClinGen Region | ISCA ID | Haploinsufficiency | Triplosensitivity | Coordinates (GRCh37/hg19) |
|---|---|---|---|---|
| 1 copy: 14q telomere; 3 copies: 14q telomere | ISCA-37477 | Haploinsufficiency unlikely | Triplosensitivity unlikely | chr14:106,050,000-107,289,540 |

| Gene | NeuroSCORE |
|---|---|
| *KIAA0125* | 0 |

| ClinGen Region | ISCA ID | Haploinsufficiency | Triplosensitivity | Coordinates (GRCh37/hg19) |
|---|---|---|---|---|
| 15q11.2 recurrent region (BP1-BP2) (includes NIPA1) | ISCA-37448 | 2 | Triplosensitivity unlikely | chr15:22,832,519-23,090,897 |

| Gene | NeuroSCORE |
|---|---|
| *TUBGCP5* | 0 |
| *CYFIP1* | 3 |
| *NIPA2* | 2 |
| *NIPA1* | 2 |

Supplementary Table 3: ClinGen Dosage Sensitivity with NeruoSCORE by Gene

| ClinGen Region | | ISCA ID | Haploinsufficiency | Triplosensitivity | Coordinates (GRCh37/hg19) |
|---|---|---|---|---|---|
| 15q11q13 recurrent (PWS/AS) region (BP1-BP3, Class 1) | | ISCA-37404 | 3 | 3 | chr15:22,832,519-28,379,874 |
| **Gene** | **NeuroSCORE** | **Gene** | **NeuroSCORE** | **Gene** | **NeuroSCORE** |
| *TUBGCP5* | 0 | *GOLGA6L2* | 0 | *UBE3A* | 4 |
| *CYFIP1* | 3 | *MKRN3* | 0 | *ATP10A* | 0 |
| *NIPA2* | 2 | *MAGEL2* | 1 | *GABRB3* | 4 |
| *NIPA1* | 2 | *NDN* | 2 | *GABRA5* | 2 |
| *GOLGA8I* | NA | *NPAP1* | 0 | *GABRG3* | 0 |
| *RP11-467N20.5* | NA | *SNRPN* | 2 | *OCA2* | 0 |
| *GOLGA8S* | 0 | *SNURF* | 1 | *HERC2* | 4 |

| ClinGen Region | | ISCA ID | Haploinsufficiency | Triplosensitivity | Coordinates (GRCh37/hg19) |
|---|---|---|---|---|---|
| 15q11q13 recurrent (PWS/AS) region (BP2-BP3, Class 2) | | ISCA-37478 | 3 | 3 | chr15:23,747,996-28,379,874 |
| **Gene** | **NeuroSCORE** | **Gene** | **NeuroSCORE** | | |
| *MKRN3* | 0 | *ATP10A* | 0 | | |
| *MAGEL2* | 1 | *GABRB3* | 4 | | |
| *NDN* | 2 | *GABRA5* | 2 | | |
| *NPAP1* | 0 | *GABRG3* | 0 | | |
| *SNRPN* | 2 | *OCA2* | 0 | | |
| *SNURF* | 1 | *HERC2* | 4 | | |
| *UBE3A* | 4 | | | | |

Supplementary Table 3: ClinGen Dosage Sensitivity with NeruoSCORE by Gene

| ClinGen Region | ISCA ID | Haploinsufficiency | Triplosensitivity | Coordinates (GRCh37/hg19) |
|---|---|---|---|---|
| 15q13 recurrent region (BP3-BP4) (includes APBA2) | ISCA-46285 | 0 | 0 | chr15:29,156,959-30,368,990 |

| Gene | NeuroSCORE |
|---|---|
| *APBA2* | 2 |
| *FAM189A1* | 1 |
| *NSMCE3* | 0 |
| *TJP1* | 2 |

| ClinGen Region | ISCA ID | Haploinsufficiency | Triplosensitivity | Coordinates (GRCh37/hg19) |
|---|---|---|---|---|
| 15q13.3 recurrent region (BP4-BP5) (includes CHRNA7) | ISCA-37411 | 3 | 1 | chr15:31,192,889-32,445,405 |

| Gene | NeuroSCORE |
|---|---|
| *FAN1* | 1 |
| *MTMR10* | 1 |
| *TRPM1* | 0 |
| *KLF13* | 3 |
| *OTUD7A* | 2 |
| *CHRNA7* | 0 |

| ClinGen Region | ISCA ID | Haploinsufficiency | Triplosensitivity | Coordinates (GRCh37/hg19) |
|---|---|---|---|---|
| 15q13.3 recurrent region (D-CHRNA7 to BP5) (includes CHRNA7 and OTUD7A) | ISCA-46295 | 3 | Triplosensitivity unlikely | chr15:32,019,621-32,445,405 |

| Gene | NeuroSCORE |
|---|---|
| *OTUD7A* | 2 |
| *CHRNA7* | 0 |

Supplementary Table 3: ClinGen Dosage Sensitivity with NeruoSCORE by Gene

| ClinGen Region | | ISCA ID | Haploinsufficiency | Triplosensitivity | Coordinates (GRCh37/hg19) |
|---|---|---|---|---|---|
| 15q24 recurrent region (A-D) (includes SIN3A) | | ISCA-37396 | 3 | 1 | chr15:72,963,715-75,972,909 |
| **Gene** | **NeuroSCORE** | **Gene** | **NeuroSCORE** | **Gene** | **NeuroSCORE** |
| *HIGD2B* | 0 | *STRA6* | 0 | *COX5A* | 2 |
| *BBS4* | 0 | *CCDC33* | 0 | *RPP25* | 0 |
| *ADPGK* | 1 | *CYP11A1* | 0 | *SCAMP5* | 2 |
| *NEO1* | 2 | *SEMA7A* | 1 | *PPCDC* | 0 |
| *HCN4* | 1 | *UBL7* | 2 | *C15orf39* | 0 |
| *C15orf60* | 0 | *ARID3B* | 0 | *GOLGA6C* | 0 |
| *NPTN* | 4 | *CLK3* | 1 | *GOLGA6D* | 0 |
| *CD276* | 0 | *EDC3* | 0 | *COMMD4* | 2 |
| *INSYN1* | 2 | *CYP1A1* | 0 | *NEIL1* | 0 |
| *TBC1D21* | 0 | *CYP1A2* | 0 | *MAN2C1* | 1 |
| *LOXL1* | 0 | *CSK* | 3 | *SIN3A* | 3 |
| *STOML1* | 1 | *LMAN1L* | 0 | *PTPN9* | 1 |
| *PML* | 1 | *CPLX3* | 0 | *SNUPN* | 1 |
| *GOLGA6* | 0 | *ULK3* | 2 | *IMP3* | 2 |
| *ISLR2* | 0 | *SCAMP2* | 1 | *SNX33* | 0 |
| *RP11-247C2.2* | 0 | *MPI* | 2 | *CSPG4* | 0 |
| *ISLR* | 0 | *FAM219B* | 2 | *AC105020.1* | 0 |

Supplementary Table 3: ClinGen Dosage Sensitivity with NeruoSCORE by Gene

| ClinGen Region | ISCA ID | Haploinsufficiency | Triplosensitivity | Coordinates (GRCh37/hg19) |
|---|---|---|---|---|
| 15q24 recurrent region (A-C) | ISCA-46296 | 3 | 1 | chr15:72,963,715-75,508,312 |

| Gene | NeuroSCORE | Gene | NeuroSCORE | Gene | NeuroSCORE |
|---|---|---|---|---|---|
| *HIGD2B* | 0 | *GOLGA6A* | 0 | *CYP1A2* | 0 |
| *BBS4* | 0 | *ISLR2* | 0 | *CSK* | 3 |
| *ADPGK* | 1 | *RP11-247C2.2* | 0 | *LMAN1L* | 0 |
| *NEO1* | 2 | *ISLR* | 0 | *CPLX3* | 0 |
| *HCN4* | 1 | *STRA6* | 0 | *ULK3* | 2 |
| *C15orf60* | 0 | *CCDC33* | 0 | *SCAMP2* | 1 |
| *NPTN* | 4 | *CYP11A1* | 0 | *MPI* | 2 |
| *CD276* | 0 | *SEMA7A* | 1 | *FAM219B* | 2 |
| *INSYN1* | 2 | *UBL7* | 2 | *COX5A* | 2 |
| *TBC1D21* | 0 | *ARID3B* | 0 | *RPP25* | 0 |
| *LOXL1* | 0 | *CLK3* | 1 | *SCAMP5* | 2 |
| *STOML1* | 1 | *EDC3* | 0 | *PPCDC* | 0 |
| *PML* | 1 | *CYP1A1* | 0 | *C15orf39* | 0 |

| ClinGen Region | ISCA ID | Haploinsufficiency | Triplosensitivity | Coordinates (GRCh37/hg19) |
|---|---|---|---|---|
| 15q24 recurrent region (C-D) (includes SIN3A) | ISCA-46300 | 3 | 0 | chr15:75,631,787-75,972,909 |

| Gene | NeuroSCORE | Gene | NeuroSCORE |
|---|---|---|---|
| *COMMD4* | 2 | *SNUPN* | 1 |
| *NEIL1* | 0 | *IMP3* | 2 |
| *MAN2C1* | 1 | *SNX33* | 0 |
| *SIN3A* | 3 | *CSPG4* | 0 |
| *PTPN9* | 1 | *AC105020.1* | 0 |

Supplementary Table 3: ClinGen Dosage Sensitivity with NeruoSCORE by Gene

| ClinGen Region | | ISCA ID | Haploinsufficiency | Triplosensitivity | Coordinates (GRCh37/hg19) |
|---|---|---|---|---|---|
| 15q25.2 recurrent region (LCR B-C, proximal) | | ISCA-37500 | 3 | 0 | chr15:83,213,988-84,714,733 |
| **Gene** | **NeuroSCORE** | **Gene** | **NeuroSCORE** | **Gene** | **NeuroSCORE** |
| *RP11-152F13.10* | 1 | *WHAMM* | 0 | *TM6SF1* | 0 |
| *RP11-379H8.1* | 0 | *HOMER2* | 0 | *HDGFL3* | 3 |
| *CPEB1* | 2 | *FAM103A1* | 0 | *BNC1* | 1 |
| *AP3B2* | 2 | *C15orf40* | 0 | *SH3GL3* | 2 |
| *FSD2* | 0 | *BTBD1* | 2 | *ADAMTSL3* | 0 |

| ClinGen Region | | ISCA ID | Haploinsufficiency | Triplosensitivity | Coordinates (GRCh37/hg19) |
|---|---|---|---|---|---|
| 15q25.2q25.3 recurrent region (LCR C-D, distal) | | ISCA-37514 | 1 | 0 | chr15:85,139,652-85,713,001 |
| **Gene** | **NeuroSCORE** | **Gene** | **NeuroSCORE** | | |
| *ZSCAN2* | 0 | *ZNF592* | 1 | | |
| *WDR73* | 0 | *ALPK30* | 0 | | |
| *NMB* | 0 | *SLC28A1* | 0 | | |
| *SEC11A* | 2 | *PDE8A* | 1 | | |

| ClinGen Region | ISCA ID | Haploinsufficiency | Triplosensitivity | Coordinates (GRCh37/hg19) |
|---|---|---|---|---|
| 3 copies: 15q telomere | ISCA-37480 | 0 | Triplosensitivity unlikely | chr15:102,161,480-102,521,392 |
| **Gene** | **NeuroSCORE** | **Gene** | **NeuroSCORE** | |
| *TM2D3* | 2 | *OR4F15* | 0 | |
| *TARSL2* | 1 | *OR4F4* | 0 | |
| *RP11-89K11.1* | 0 | | | |
| *OR4F6* | 0 | | | |

Supplementary Table 3: ClinGen Dosage Sensitivity with NeruoSCORE by Gene

| ClinGen Region | ISCA ID | Haploinsufficiency | Triplosensitivity | Coordinates (GRCh37/hg19) |
|---|---|---|---|---|
| 16p13.3 region (includes CREBBP) | ISCA-37406 | 3 | 1 | chr16:3,775,056-3,930,121 |

| Gene | NeuroSCORE |
|---|---|
| *CREBBP* | 3 |

| ClinGen Region | ISCA ID | Haploinsufficiency | Triplosensitivity | Coordinates (GRCh37/hg19) |
|---|---|---|---|---|
| 16p13.11 recurrent region (BP2-BP3) (includes MYH11) | ISCA-37415 | 3 | 2 | chr16:15,511,711-16,292,265 |

| Gene | NeuroSCORE | Gene | NeuroSCORE |
|---|---|---|---|
| *RP11-1021N1.1* | NA | *MYH11* | 3 |
| *C16orf45* | 2 | *FOPNL* | 1 |
| *MARF1* | 4 | *ABCC1* | 0 |
| *NDE1* | 0 | *ABCC6* | 0 |

| ClinGen Region | ISCA ID | Haploinsufficiency | Triplosensitivity | Coordinates (GRCh37/hg19) |
|---|---|---|---|---|
| 16p12.2 recurrent region (includes OTOA) (distal region) | ISCA-46297 | Gene associated with AR phenotype | Triplosensitivity unlikely | chr16:21,570,113-21,740,423 |

| Gene | NeuroSCORE |
|---|---|
| *METTL9* | 2 |
| *IGSF6* | 0 |
| *OTOA* | 0 |

Supplementary Table 3: ClinGen Dosage Sensitivity with NeruoSCORE by Gene

| ClinGen Region | | ISCA ID | Haploinsufficiency | Triplosensitivity | Coordinates (GRCh37/hg19) |
|---|---|---|---|---|---|
| 16p12.2 recurrent region (includes EEF2K, CDR2) (proximal region) | | ISCA-37409 | 2 | 0 | chr16:21,948,445-22,430,804 |
| **Gene** | **NeuroSCORE** | **Gene** | **NeuroSCORE** | | |
| *UQCRC2* | 2 | *SDR42E2* | 0 | | |
| *PDZD9* | 0 | *EEF2K* | 0 | | |
| *MOSMO* | 0 | *POLR3E* | 0 | | |
| *VWA3A* | 0 | *CDR2* | 1 | | |

| ClinGen Region | | ISCA ID | Haploinsufficiency | Triplosensitivity | Coordinates (GRCh37/hg19) |
|---|---|---|---|---|---|
| 16p11.2 recurrent region (distal, BP2-BP3) (includes SH2B1) | | ISCA-37486 | 3 | 1 | chr16:28,822,635-29,046,499 |
| **Gene** | **NeuroSCORE** | **Gene** | **NeuroSCORE** | | |
| *ATXN2L* | 4 | *CD19* | 0 | | |
| *TUFM* | 2 | *NFATC2IP* | 0 | | |
| *SH2B1* | 4 | *SPNS1* | 2 | | |
| *ATP2A1* | 0 | *LAT* | 0 | | |
| *RABEP2* | 0 | | | | |

Supplementary Table 3: ClinGen Dosage Sensitivity with NeruoSCORE by Gene

| ClinGen Region | | ISCA ID | Haploinsufficiency | Triplosensitivity | Coordinates (GRCh37/hg19) |
|---|---|---|---|---|---|
| 16p11.2 recurrent region (proximal, BP4-BP5) (includes TBX6) | | [ISCA-37400](ISCA-37400) | 3 | 3 | chr16:29,649,997-30,199,852 |
| **Gene** | **NeuroSCORE** | **Gene** | **NeuroSCORE** | | |
| *SPN* | 0 | *TAOK2* | 4 | | |
| *QPRT* | 1 | *HIRIP3* | 1 | | |
| *C16orf54* | 0 | *INO80E* | 2 | | |
| *ZG16* | 0 | *DOC2A* | 2 | | |
| *KIF22* | 0 | *C16orf92* | 1 | | |
| *MAZ* | 2 | *FAM57B* | 1 | | |
| *PRRT2* | 2 | *ALDOA* | 2 | | |
| *PAGR1* | 2 | *PPP4C* | 3 | | |
| *MVP* | 1 | *TBX6* | 0 | | |
| *CDIPT* | 2 | *YPEL3* | 2 | | |
| *SEZ6L2* | 2 | *GDPD3* | 0 | | |
| *ASPHD1* | 2 | *MAPK3* | 2 | | |
| *KCTD13* | 2 | *CORO1A* | 3 | | |
| *TMEM219* | 2 | | | | |

| ClinGen Region | ISCA ID | Haploinsufficiency | Triplosensitivity | Coordinates (GRCh37/hg19) |
|---|---|---|---|---|
| 3 copies: 16p centromere | ISCA-37481 | Haploinsufficiency unlikely | Triplosensitivity unlikely | chr16:34,202,088-35,147,508 |
| **Gene** | **NeuroSCORE** | | | |
| *CTD-2144E22.5* | 0 | | | |

Supplementary Table 3: ClinGen Dosage Sensitivity with NeruoSCORE by Gene

| ClinGen Region | | ISCA ID | Haploinsufficiency | Triplosensitivity | Coordinates (GRCh37/hg19) |
|---|---|---|---|---|---|
| 17p13.3 (Miller-Dieker syndrome) region (includes YWHAE and PAFAH1B1) | | ISCA-37430 | 3 | 3 | chr17:1,247,833-2,588,909 |
| **Gene** | **NeuroSCORE** | **Gene** | **NeuroSCORE** | | |
| *YWHAE* | 4 | *RPA1* | 2 | | |
| *CRK* | 3 | *RTN4RL1* | 1 | | |
| *MYO1C* | 0 | *DPH1* | 2 | | |
| *INPP5K* | 2 | *OVCA2* | NA | | |
| *PITPNA* | 3 | *HIC1* | 1 | | |
| *SLC43A2* | 1 | *SMG6* | 1 | | |
| *SCARF1* | 0 | *SRR* | 1 | | |
| *RILP* | 0 | *TSR1* | 1 | | |
| *PRPF8* | 4 | *SGSM2* | 2 | | |
| *TLCD2* | 0 | *MNT* | 1 | | |
| *WDR81* | 1 | *METTL16* | 0 | | |
| *SERPINF2* | 0 | *AC006435.1* | NA | | |
| *SERPINF1* | 0 | *PAFAH1B1* | 4 | | |
| *SMYD4* | 0 | | | | |

| ClinGen Region | | ISCA ID | Haploinsufficiency | Triplosensitivity | Coordinates (GRCh37/hg19) |
|---|---|---|---|---|---|
| 17p12 recurrent (HNPP/CMT1A) region (includes PMP22) | | ISCA-37436 | 3 | 3 | chr17:14,097,915-15,422,952 |
| **Gene** | **NeuroSCORE** | **Gene** | **NeuroSCORE** | | |
| *COX10* | 0 | *TEKT3* | 0 | | |
| *CDRT15* | 0 | *CDRT4* | 1 | | |
| *HS3ST3B1* | 0 | *TVP23C-CDRT4* | NA | | |
| *PMP22* | 1 | *TVP23C* | 0 | | |

Supplementary Table 3: ClinGen Dosage Sensitivity with NeruoSCORE by Gene

| ClinGen Region | | ISCA ID | Haploinsufficiency | Triplosensitivity | Coordinates (GRCh37/hg19) |
|---|---|---|---|---|---|
| 17p11.2 recurrent (SMS/PLS) region (includes RAI1) | | ISCA-37418 | 3 | 3 | chr17:16,810,028-20,213,202 |
| **Gene** | **NeuroSCORE** | **Gene** | **NeuroSCORE** | **Gene** | **NeuroSCORE** |
| *TNFRSF13B* | 0 | *ALKBH5* | 2 | *FAM83G* | 0 |
| *MPRIP* | 3 | *LLGL1* | 2 | *GRAP* | 0 |
| *PLD6* | 0 | *FLII* | 2 | *AC007952.5* | 0 |
| *FLCN* | 1 | *MIEF2* | 0 | *GRAPL* | 0 |
| *COPS3* | 3 | *TOP3A* | 0 | *EPN2* | 2 |
| *NT5M* | 0 | *SMCR8* | 0 | *B9D1* | 0 |
| *MED9* | 1 | *SHMT1* | 0 | *MAPK7* | 0 |
| *RASD1* | 1 | *EVPLL* | 0 | *MAPK4* | 0 |
| *PEMT* | 2 | *LGALS9C* | 0 | *RNF112* | 2 |
| *RAI1* | 1 | *FAM106A* | 0 | *SLC47A1* | 0 |
| *SREBF1* | 1 | *TBC1D28* | 0 | *ALDH3A2* | 2 |
| *TOM1L2* | 1 | *ZNF286B* | 0 | *SLC47A2* | 0 |
| *DRC3* | 0 | *TRIM16L* | 0 | *ALDH3A1* | 0 |
| *ATPAF2* | 0 | *FBXW10* | 0 | *ULK2* | 3 |
| *GID4* | 0 | *TVP23B* | 0 | *AKAP10* | 1 |
| *DRG2* | 2 | *PRPSAP2* | 2 | *SPECC1* | 0 |
| *MYO15A* | 0 | *SLC5A10* | 1 | | |

Supplementary Table 3: ClinGen Dosage Sensitivity with NeruoSCORE by Gene

| ClinGen Region | ISCA ID | Haploinsufficiency | Triplosensitivity | Coordinates (GRCh37/hg19) |
|---|---|---|---|---|
| 17q11.2 recurrent region (includes NF1) | ISCA-37431 | 3 | 2 | chr17:29,097,069-30,264,027 |

| Gene | NeuroSCORE | Gene | NeuroSCORE |
|---|---|---|---|
| *CRLF3* | 0 | *CTD-2370N5.3* | NA |
| *ATAD5* | 1 | *EVI2A* | 1 |
| *TEFM* | 0 | *RAB11FIP4* | 4 |
| *ADAP2* | 0 | *AC003101.1* | 0 |
| *RNF135* | 0 | *NRBF2* | 2 |
| *NF1* | 3 | *COPRS* | 2 |
| *OMG* | 2 | *UTP6* | 0 |
| *EVI2B* | 0 | | |

| ClinGen Region | ISCA ID | Haploinsufficiency | Triplosensitivity | Coordinates (GRCh37/hg19) |
|---|---|---|---|---|
| 17q12 recurrent (RCAD syndrome) region (includes HNF1B) | ISCA-37432 | 3 | 3 | chr17:34,815,072-36,192,489 |

| Gene | NeuroSCORE | Gene | NeuroSCORE | Gene | NeuroSCORE |
|---|---|---|---|---|---|
| *ZNHIT3* | 2 | *MRM1* | 0 | *TADA2A* | 0 |
| *MYO19* | 0 | *LHX1* | 1 | *DUSP14* | 2 |
| *PIGW* | 0 | *AATF* | 2 | *SYNRG* | 2 |
| *GGNBP2* | 4 | *ACACA* | 5 | *DDX52* | 0 |
| *DHRS11* | 1 | *C17orf78* | 0 | *HNF1B* | 2 |

Supplementary Table 3: ClinGen Dosage Sensitivity with NeruoSCORE by Gene

| ClinGen Region | ISCA ID | Haploinsufficiency | Triplosensitivity | Coordinates (GRCh37/hg19) |
|---|---|---|---|---|
| 17q21.3 recurrent region (includes KANSL1) | ISCA-37420 | 3 | 1 | chr17:43,705,166-44,164,880 |

| Gene | NeuroSCORE |
|---|---|
| *CRHR1* | 0 |
| *SPPL2C* | 0 |
| *MAPT* | 3 |
| *STH* | 0 |
| *KANSL1* | 2 |

| ClinGen Region | ISCA ID | Haploinsufficiency | Triplosensitivity | Coordinates (GRCh37/hg19) |
|---|---|---|---|---|
| 17q23.1q23.2 recurrent region (includes TBX2, TBX4) | ISCA-37501 | 3 | 2 | chr17:58,113,002-60,275,809 |

| Gene | NeuroSCORE | Gene | NeuroSCORE | Gene | NeuroSCORE |
|---|---|---|---|---|---|
| *HEATR6* | 0 | *RP11-15E18.4* | 0 | *TBX4* | 0 |
| *CA4* | 0 | *PPM1D* | 0 | *NACA2* | 1 |
| *USP32* | 2 | *BCAS3* | 1 | *BRIP1* | 0 |
| *C17orf64* | 0 | *TBX2* | 1 | *INTS2* | 1 |
| *APPBP2* | 1 | *C17orf82* | 0 | *MED13* | 1 |

| ClinGen Region | ISCA ID | Haploinsufficiency | Triplosensitivity | Coordinates (GRCh37/hg19) |
|---|---|---|---|---|
| 19q13.3 region (PSG gene cluster) | ISCA-37483 | Haploinsufficiency unlikely | Triplosensitivity unlikely | chr19:43,242,796-43,741,310 |

| Gene | NeuroSCORE | Gene | NeuroSCORE |
|---|---|---|---|
| *PSG3* | 0 | *PSG11* | 0 |
| *PSG8* | 0 | *PSG2* | 0 |
| *PSG1* | 0 | *PSG5* | 0 |
| *PSG6* | 0 | *PSG4* | 0 |
| | | *PSG9* | 0 |

Supplementary Table 3: ClinGen Dosage Sensitivity with NeruoSCORE by Gene

| ClinGen Region | | ISCA ID | Haploinsufficiency | Triplosensitivity | Coordinates (GRCh37/hg19) |
|---|---|---|---|---|---|
| 22q11.21 recurrent (Cat eye syndrome) region (includes CECR2) | | ISCA-37393 | 0 | 3 | chr22:17,392,953-18,591,860 |
| **Gene** | **NeuroSCORE** | **Gene** | **NeuroSCORE** | | |
| *GAB4* | NA | *SLC25A18* | 1 | | |
| *IL17RA* | 0 | *ATP6V1E1* | 2 | | |
| *CECR6* | 0 | *BCL2L13* | 0 | | |
| *AC006946.15* | 0 | *BID* | 2 | | |
| *CECR5* | 2 | *MICAL3* | 1 | | |
| *ADA2* | 0 | *PEX26* | 1 | | |
| *CECR2* | 1 | | | | |

| ClinGen Region | | ISCA ID | Haploinsufficiency | Triplosensitivity | Coordinates (GRCh37/hg19) |
|---|---|---|---|---|---|
| 22q11.2 recurrent (DGS/VCFS) region (proximal, A-D) (includes TBX1) | | ISCA-37446 | 3 | 3 | chr22:18,912,231-21,465,672 |
| **Gene** | **NeuroSCORE** | **Gene** | **NeuroSCORE** | **Gene** | **NeuroSCORE** |
| *PRODH* | 1 | *TBX1* | 0 | *FAM230A* | 0 |
| *DGCR2* | 2 | *GNB1L* | 0 | *USP41* | 0 |
| *ESS2* | 0 | *C22orf29* | 0 | *ZNF74* | 0 |
| *TSSK2* | NA | *TXNRD2* | 0 | *SCARF2* | 1 |
| *GSC2* | 0 | *COMT* | 2 | *KLHL22* | 2 |
| *SLC25A1* | 2 | *ARVCF* | 1 | *MED15* | 4 |
| *CLTCL1* | 0 | *TANGO2* | 0 | *PI4KA* | 2 |
| *HIRA* | 3 | *DGCR8* | 1 | *SERPIND1* | 0 |
| *C22orf39* | 2 | *TRMT2A* | 1 | *SNAP29* | 1 |
| *MRPL40* | 2 | *RANBP1* | 2 | *CRKL* | 2 |
| *UFD1* | 3 | *ZDHHC8* | 1 | *AIFM3* | 2 |
| *CDC45* | 0 | *CCDC188* | 0 | *LZTR1* | 1 |
| *CLDN5* | 3 | *RTN4R* | 2 | *THAP7* | 1 |
| *SEPTIN5* | 1 | *DGCR6L* | 2 | *P2RX6* | 0 |
| *GP1BB* | 1 | *RIMBP3* | 0 | *SLC7A4* | 0 |
| | | | | *LRRC74B* | 0 |

Supplementary Table 3: ClinGen Dosage Sensitivity with NeruoSCORE by Gene

| ClinGen Region | ISCA ID | Haploinsufficiency | Triplosensitivity | Coordinates (GRCh37/hg19) |
|---|---|---|---|---|
| 22q11.2 recurrent (DGS/VCFS) region (proximal, A-B) (includes TBX1) | ISCA-37433 | 3 | 3 | chr22:18,912,231-20,287,208 |

| Gene | NeuroSCORE | Gene | NeuroSCORE |
|---|---|---|---|
| *PRODH* | 1 | *GP1BB* | 1 |
| *DGCR2* | 2 | *TBX1* | 0 |
| *ESS2* | 0 | *GNB1L* | 0 |
| *TSSK2* | NA | *C22orf29* | 0 |
| *GSC2* | 0 | *TXNRD2* | 0 |
| *SLC25A1* | 2 | *COMT* | 2 |
| *CLTCL1* | 0 | *ARVCF* | 1 |
| *HIRA* | 3 | *TANGO2* | 0 |
| *C22orf39* | 2 | *DGCR8* | 1 |
| *MRPL40* | 2 | *TRMT2A* | 1 |
| *UFD1* | 3 | *RANBP1* | 2 |
| *CDC45* | 0 | *ZDHHC8* | 1 |
| *CLDN5* | 3 | *CCDC188* | 0 |
| *SEPTIN5* | 1 | *RTN4R* | 2 |

Supplementary Table 3: ClinGen Dosage Sensitivity with NeruoSCORE by Gene

| ClinGen Region | | ISCA ID | Haploinsufficiency | Triplosensitivity | Coordinates (GRCh37/hg19) |
|---|---|---|---|---|---|
| 22q11.2 recurrent region (central, B/C-D) (includes CRKL) | | ISCA-37516 | 2 | 1 | chr22:20,731,986-21,465,672 |
| **Gene** | **NeuroSCORE** | **Gene** | **NeuroSCORE** | | |
| *USP41* | 0 | *CRKL* | 2 | | |
| *ZNF74* | 0 | *AIFM3* | 2 | | |
| *SCARF2* | 1 | *LZTR1* | 1 | | |
| *KLHL22* | 2 | *THAP7* | 1 | | |
| *MED15* | 4 | *P2RX6* | 0 | | |
| *PI4KA* | 2 | *SLC7A4* | 0 | | |
| *SERPIND1* | 0 | *LRRC74B* | 0 | | |
| *SNAP29* | 1 | | | | |

| ClinGen Region | | ISCA ID | Haploinsufficiency | Triplosensitivity | Coordinates (GRCh37/hg19) |
|---|---|---|---|---|---|
| 22q11.2 recurrent region (distal type I, D-E/F) | | ISCA-37397 | 3 | 3 | chr22:21,917,117-23,649,111 |
| **Gene** | **NeuroSCORE** | **Gene** | **NeuroSCORE** | | |
| *UBE2L3* | 3 | *ZNF280B* | 0 | | |
| *YDJC* | 1 | *ZNF280A* | 0 | | |
| *CCDC116* | 0 | *PRAME* | 0 | | |
| *SDF2L1* | 1 | *LL22NC03-63E9.3* | 0 | | |
| *PPIL2* | 1 | *GGTLC2* | 0 | | |
| *YPEL1* | 1 | *IGLL5* | NA | | |
| *MAPK1* | 4 | *RSPH14* | 1 | | |
| *PPM1F* | 1 | *GNAZ* | 3 | | |
| *TOP3B* | 0 | *RAB36* | 0 | | |
| *VPREB1* | 0 | *BCR* | 3 | | |

Supplementary Table 3: ClinGen Dosage Sensitivity with NeruoSCORE by Gene

| ClinGen Region | ISCA ID | Haploinsufficiency | Triplosensitivity | Coordinates (GRCh37/hg19) |
|---|---|---|---|---|
| Xp22.31 recurrent region (includes STS) | ISCA-37417 | 3 | Triplosensitivity unlikely | chrX:6,455,812-8,124,954 |
| **Gene** | **NeuroSCORE** | | | |
| *PUDP* | 0 | | | |
| *STS* | 0 | | | |
| *VCX* | 0 | | | |
| *PNPLA4* | 0 | | | |

| ClinGen Region | ISCA ID | Haploinsufficiency | Triplosensitivity | Coordinates (GRCh37/hg19) |
|---|---|---|---|---|
| Xp21.2 region (includes NR0B1) | ISCA-46302 | 0 | 3 | chrX:30,195,000-30,355,000 |
| **Gene** | **NeuroSCORE** | | | |
| *NR0B1* | 1 | | | |
| *MAGEB1* | 0 | | | |
| *MAGEB2* | 0 | | | |
| *MAGEB3* | 0 | | | |
| *MAGEB4* | 0 | | | |

| ClinGen Region | ISCA ID | Haploinsufficiency | Triplosensitivity | Coordinates (GRCh37/hg19) |
|---|---|---|---|---|
| Xp11.23 region (includes MAOA and MAOB) | ISCA-37468 | 3 | 0 | chrX:43,514,154-43,741,720 |
| **Gene** | **NeuroSCORE** | | | |
| *MAOA* | 2 | | | |
| *MAOB* | 2 | | | |

Supplementary Table 3: ClinGen Dosage Sensitivity with NeruoSCORE by Gene

| ClinGen Region | | ISCA ID | Haploinsufficiency | Triplosensitivity | Coordinates (GRCh37/hg19) |
|---|---|---|---|---|---|
| Xp11.22p11.23 recurrent region (includes SHROOM4) | | ISCA-46290 | 0 | 3 | chrX:48,306,152-52,103,258 |
| **Gene** | **NeuroSCORE** | **Gene** | **NeuroSCORE** | **Gene** | **NeuroSCORE** |
| *SLC38A5* | 3 | *KCND1* | 0 | *GAGE2C* | NA |
| *FTSJ1* | 3 | *GRIPAP1* | 3 | *GAGE2B* | NA |
| *PORCN* | 2 | *TFE3* | 3 | *GAGE12H* | 0 |
| *EBP* | 2 | *CCDC120* | 0 | *GAGE2A* | 0 |
| *TBC1D25* | 0 | *PRAF2* | 2 | *GAGE1* | 0 |
| *AC11568.1* | 0 | *AF196779.12* | 1 | *PAGE1* | 0 |
| *RBM3* | 2 | *WDR45* | 4 | *PAGE4* | 0 |
| *WDR13* | 2 | *GPKOW* | 3 | *USP27X* | 0 |
| *WAS* | 1 | *MAGIX* | 0 | *CLCN5* | 1 |
| *SUV39H1* | 1 | *PLP2* | 0 | *AKAP4* | 1 |
| *GLOD5* | 0 | *PRICKLES3* | 0 | *CCNB3* | 0 |
| *GATA1* | 1 | *SYP* | 2 | *GDKK* | NA |
| *HDAC6* | 3 | *CACNA1F* | 0 | *SHROOM4* | 1 |
| *ERAS* | 0 | *CCDC22* | 3 | *BMP15* | 0 |
| *PCSK1N* | 2 | *FOXP3* | 1 | *NUDT10* | 0 |
| *TIMM17B* | 2 | *PPP1R3F* | 1 | *CXorf67* | 0 |
| *PQBP1* | 3 | *GAGE10* | 0 | *NUDT11* | 1 |
| *SLC35A2* | 1 | *GAGE12J* | 0 | *GSPT2* | 1 |
| *PIM2* | 1 | *GAGE13* | 0 | *MAGED1* | 3 |
| *OTUD5* | 4 | *GAGE2D* | NA | *MAGED4* | 0 |

Supplementary Table 3: ClinGen Dosage Sensitivity with NeruoSCORE by Gene

| ClinGen Region | ISCA ID | Haploinsufficiency | Triplosensitivity | Coordinates (GRCh37/hg19) |
|---|---|---|---|---|
| Xp11.22 region (includes HUWE1) | ISCA-46299 | 0 | 3 | chrX:53,363,456-53,793,054 |

| Gene | NeuroSCORE |
|---|---|
| *SMC1A* | 3 |
| *RIBC1* | 1 |
| *HSD17B10* | 2 |
| *HUWE1* | 4 |

| ClinGen Region | ISCA ID | Haploinsufficiency | Triplosensitivity | Coordinates (GRCh37/hg19) |
|---|---|---|---|---|
| Xq28 recurrent region (includes GDI1) | ISCA-37439 | 0 | 3 | chrX:153,624,564-153,783,898 |

| Gene | NeuroSCORE | Gene | NeuroSCORE |
|---|---|---|---|
| *RPL10* | 2 | *LAGE3* | 2 |
| *DNASE1L1* | 0 | *UBL4A* | 2 |
| *TAZ* | 1 | *SLC10A3* | 0 |
| *ATP6AP1* | 3 | *FAM3A* | 2 |
| *GDI1* | 3 | *G6PD* | 3 |
| *FAM50A* | 3 | *IKBKG* | 0 |
| *PLXNA3* | 0 | | |

| ClinGen Region | ISCA ID | Haploinsufficiency | Triplosensitivity | Coordinates (GRCh37/hg19) |
|---|---|---|---|---|
| Xq28 recurrent region (int22h1/int22h2-flanked) (includes RAB39B) | ISCA-37494 | 3 | 3 | chrX:154,118,603-154,564,401 |

| Gene | NeuroSCORE | Gene | NeuroSCORE |
|---|---|---|---|
| *F8* | 1 | *BRCC3* | 0 |
| *FUNDC2* | 1 | *VBP1* | 2 |
| *CMC4* | NA | *RAB39B* | 0 |
| *MTCP1* | 0 | *CLIC2* | 0 |

Supplementary Table 4: High Scoring Genes Associated with Non-CNS Phenotypes in OMIM

| NeuroSCORE | Genes |
|---|---|
| 5 | *ANK2, MYH9, WNK1* |
| 4 | *ACTN4, ADD1, ATP2B2, BAP1, BMPR2, CALM1, CHMP4B, CNBP, CTNNA1, CTNND1, CUL3, DCAF8, DYRK1B, FN1, FXR1, GANAB, GNAS, JAK1, MEF2A, MYO9B, PICALM, PRPF8, PSMA6, RASA1, SF3B1, SF3B4, SNRNP200, STAT1, STK11, TOP1* |
| 3 | *ACVR1B, ADCY1, ATG16L1, ATP1B1, ATP2C1, BCR, CALM2, CALM3, CFL2, COL6A1, COPA, CTSB, CYLD, DKC1, EGLN1, EHBP1, EWSR1, EXOC6B, FHL1, G6PD, GDF11, GPRASP2, HMGA1, HMGCR, HNRNPDL, IFNGR2, IKZF1, INF2, IRF2BP2, IRS2, KCNH2, KIF1B, KIF21A, LRP1, LTBP3, MAGED2, MAPK8IP1, MAPRE2, MECOM, MEN1, MFN2, MLH1, MORC2, MSN, MYH11, NCSTN, NFKBIA, NLRP1, NPM1, NUMA1 ,P4HB, PABPN1, PAX5, PIP5K1C, PKD1, PLS3, PRCC, PRG4, PRKAG2, PRPF3, PRPF31, PRPF6, PTPRF, PTPRZ1, RAB7A, RB1CC1, RELA, RIMS1, RPL11, RPL13, RPL15 ,RPL18, RPL5, RPS10, RPS15A, RPSA, SEPTIN9, SERPING1, SORT1, SPRY2, STAT3, STAT5B, SUMO1, TAB2, TCF7L2, TFE3, TNPO3, TPM3, TRAF3, TSG101, WARS* |

OMIM database accessed December 31st, 2020
